# LENG8 regulation of mRNA processing, is responsible for the control of mitochondrial activity

**DOI:** 10.1101/2021.07.17.452750

**Authors:** Yongxu Zhao, Xiaoting Wang, Yuenan Liu, Niannian Li, Shengming Wang, Zhigang Sun, Zhenfei Gao, Xiaoxu Zhang, Linfei Mao, Ru Tang, Wenyue Xue, Chunyan Li, Jian Guan, Hongliang Yi, Nan Zhang, Qiurong Ding, Feng Liu

## Abstract

The processing of mRNA is essential for the maintenance of cellular and tissue homeostasis. However, the precise regulation of this process in mammalian cells, remains largely unknown. Here we have found that LENG8 represents the mammalian orthologue of the yeast mRNA processing factor Thp3 and Sac3. We go on to demonstrate that LENG8 binds to mRNAs, associates with components of mRNA processing machinery (the TREX complex) and contributes to mRNA nuclear export to the cytoplasm. Loss of *LENG8*, leads to aberrant accumulation of poly (A)^+^ RNA in the nucleus, in both Hela cells and murine fibroblasts. Furthermore, the precipitation of LENG8, is associated with an enrichment of both mRNAs and lncRNAs, and approximately half of these are also bound by the TREX component, THOC1. However, LENG8 preferentially binds mRNAs encoding for mitochondrial proteins and depletion of this processing factor, causes a dramatic breakdown in mitochondrial ultrastructure and a reduction in mitochondrial respiratory activity. Conditional deletion of *Leng8* in mouse adipose tissues lead to a decreased body weight, and increased adipose thermogenesis. Our work has found an evolutionarily conserved mRNA processing factor that can control mitochondrial activity.

## INTRODUCTION

In eukaryotes the flow of genetic information from DNA to protein, requires the correct coupling of RNA transcription and subsequent mRNA processing. These processes include 5’end capping and splicing, 3’end cleavage, polyadenylation, and finally RNA export. As the nascent pre-mRNA emerges from the RNA polymerase II (RNAPII) complex, it is packaged as a messenger ribonucleoparticle (mRNP), the optimal configuration of which, is critical for normal pre-mRNA processing and export (**Wickramasinghe and Laskey, 2015; Lee et al., 2013**). The biogenesis of mRNPs is tightly regulated by the THO complex, which has been found to be highly conserved from yeast to mammals (**Katahira and Yoneda, 2009; Yuan et al., 2018**).

In yeast, the THO complex is composed of four tightly interacting subunits, Hpr1, Tho2, Mft1, and Thp2. Numerous studies have found a relationship between THO and mRNA processing, as mutations or deletions of either subunit, caused aberrant mRNA export (**Jimeno et al., 2002; Strässer et al., 2002**). The core of the THO complex, has been found to physically associate with two additional export factors, Aly/Yra1 and RNA-dependent ATPase Sub2/UAP56. When these are also recruited to the nascent mRNA, the larger RNA-protein complex, transcription-export (TREX), is formed (**Xie & Ren, 2011**; **Zenklusen et al., 2001**).

Another protein complex, termed TREX-2, which is downstream of TREX and composed of Thp1, Sac3, Sus1, and Cdc31, is also involved in the regulation of mRNA processing (**Stewart, 2019; García-Oliver et al., 2012**). As seen with the THO complex, in yeast, loss of TREX-2 components leads to defects in mRNA export. Unlike the THO complex, which is primarily associated with active chromatin, the TREX-2 complex represents a relatively downstream step in mRNA processing. It is located primarily at the nuclear periphery, in association with the nuclear pore complex. Using a combination of biochemical and genetic analyses, several additional proteins have been identified, as mediators of mRNA processing, including Mex67 and Mtr2, which form chemical bridges with the THO and TREX-2 complexes (**Rondón et al., 2010**).

Because of their fundamental and essential roles in gene expression, both the TREX and TREX-2 complexes are evolutionarily conserved, from budding yeast to humans (**Domínguez-Sánchez et al., 2011; Cheng et al., 2006**). The human orthologues of the THO subunits, Hpr1, Tho2 and Tex1 are termed THOC1, THOC2 and THOC3, respectively (**Li et al., 2005**; **Masuda et al., 2005; Kumar et al., 2015**). However, human THO has three additional subunits, THOC5, THOC6 and THOC7 not found in yeast (**Pierce et al. 2008; Tran et al., 2014; Saran et al., 2016**). Furthermore, as found with yeast Yra1, the human homologs THOC4/ALYREF, also play essential roles in the regulation of mRNA maturation (**Chi et al., 2013; Shi et al., 2017; Fan et al., 2019**). However how mRNA processing is controlled in mammalian cells, still remains largely nebulous.

The super-helical PCI domain, which is firstly found in and named for multi-subunit complexes proteasome, CSN and eIF3, serves as the principal scaffold for Thp1-Sac3 duo in the TREX-2 complex(**Khoshnevis et al., 2014; Kragelund et al., 2016**). The Thp3-Csn12 minicomplex, which is also identified as a PCI complex, has been found to regulate transcriptional elongation and mRNA maturation in yeast (**Jimeno et al., 2011; Kragelund et al., 2016**), but its precise mechanism and the potential existence of homologous proteins in mammals, remains unknown. Using bioinformatic alignment tools, we have identified Leukocyte Receptor Cluster Member 8 (LENG8) as the mammalian homologue of Thp3 and found that it can associate with PCI domain containing 2 (PCID2), the mammalian equivalent of Csn12. LENG8 can bind mRNAs associate with the mRNA processing machinery, causing the attenuation of mRNA export. It preferentially binds to mRNAs encoding for proteins that localize to the mitochondria and its activity is required for the maintenance of mitochondrial morphology and respiratory function. This work has revealed evolutionarily conserved mRNA processing machinery, that can control mitochondrial activity.

## RESULTS

### Identification of LENG8 and PCID2 as the mammalian orthologues of the yeast Thp3-Csn12 complex

To further understand the process of mRNP biogenesis, bioinformatics analyses were performed on the yeast mRNA maturation factors and their mammalian orthologues. (**Fig. 1a; Supplementary Table 1**). The yeast Thp3-Csn12 complex, is a newly identified PCI minicomplex in yeast, the activity of which, is required for transcriptional elongation and mRNA processing. However, its exact role, and mammalian equivalents, involved in mRNP biogenesis, remains unknown. PCID2, which was previously identified as the mammalian homologue of THP1 subunit of the yeast TREX2 complex, exhibited greater similarity to CSN12, which also contained a typical PCI domain (**Bhatia et al., 2014**) (**Fig. 1a, 1b & 1d** & **Supplementary Table 1, Supplementary Fig. 1**). Furthermore, using bioinformatic alignment tools(msa R package)(**U. Bodenhofer *et al*., 2015**), we identified LENG8, CG6700 and the Hypersensitive to Pore-forming toxin (HPO-10) genes as orthologues of THP3, in mammals, drosophila and nematodes, respectively (**Fig. 1a-c & Supplementary Table 1, Supplementary Fig. 2**). Human LENG8 protein, consists of three distinct domains: the N-terminal topoisomerase II-associated protein (PAT1) region, mid-HJURP (Holliday junction recognition protein)-associated repeat domain and the C-terminal atypical SAC3-subytpe of PCI domain. All orthologues of THP3 contain this subtype of PCI domain and using algorithmic analysis, it has been found that the amino acid sequence at the PCI domain, displays 71% similarity between THP3 and LENG8, across all eukaryotes (**Fig. 1e**). Similar to the PCI domain in SAC3 and THP3, the LENG8 equivalent consists of curved helical repeats, terminating in a globular winged helix(WH) subdomain(**Fig. 1f**). Taken together, these results suggested that eukaryotic LENG8 is highly conserved.

**Figure 1.**
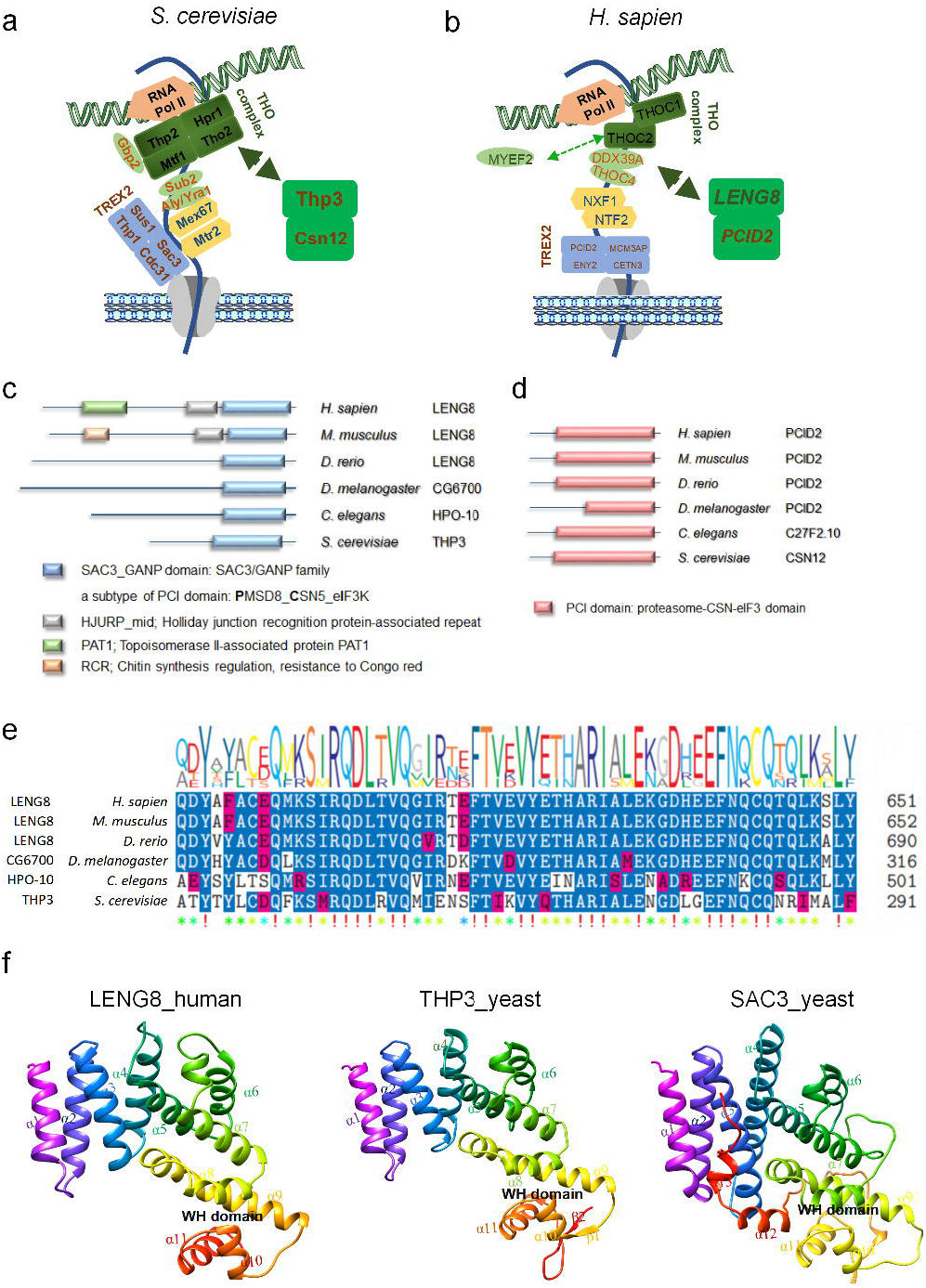
Conservation of mRNP biogenesis machinery from yeast to human. (**a**)mRNP biogenesis machinery in yeast and human. (**b**) Alignment of human LENG8 with its orthologues in other species. (**c**) Alignment of the amino acids sequence of the CPE domain.(**d**)alignment of human PCID2 with its orthologues in other species.(**e**) Alignment of the amino acids sequence of the PCI domain. (**f**) 3D structures of PCI domains from LENG8, THP3 and SAC3 proteins. Colour coding represents the identical or similar amino acids(**e**).

We also found that the ectopically expressed LENG8, as well as the endogenous LENG8, are predominantly located in the cell nucleus, especially nucleosomes and nuclear speckles (**Supplementary Fig. 3a and b**). However, a truncated mutant form of LENG8 (1-556), lacking the PCI domain, was found to locate to the cytosol, indicating that this domain is crucial for the correct nuclear localization of LENG8 (**Supplementary Fig. 3c and d**). However, the true functional relevance of this domain in LENG8, remains unknown.

Here, we hypothesized that LENG8 associates with PCID2 and can contribute to mRNA export. To test this, FLAG-tagged LENG8 was co-expressed with MYC-tagged PCID2, in HEK293T cells, and co-immunoprecipitation (coIP) experiments were performed to detect their potential interaction. Results from these experiments revealed that PCID2, was indeed detected in the pellet fraction of FLAG-tagged LENG8 (**Fig. 2a**). Furthermore, the poly(A)^+^ RNP complex, was purified using biotinylated oligo-dT and then precipitated with streptavidin conjugated beads. The mRNP complex was then subjected to Western blot analysis and the results demonstrated that LENG8, PCID2, as well as THOC1 and ALYREF, (two of the dominant components of THO), were detected in the palleted fraction of the mRNP (**Fig. 2b**). We next performed fluorescence *in situ* hybridization (FISH) analysis of the poly(A)^+^ mRNA using Cy3-labeled oligo-dT and found that LENG8 colocalized with PCID2, and that both co-localized with poly(A)^+^ RNAs (**Fig. 2c-e**). Taken together these results confirm that both LENG8 and PCID2 are associated with RNAs.

**Figure 2.**
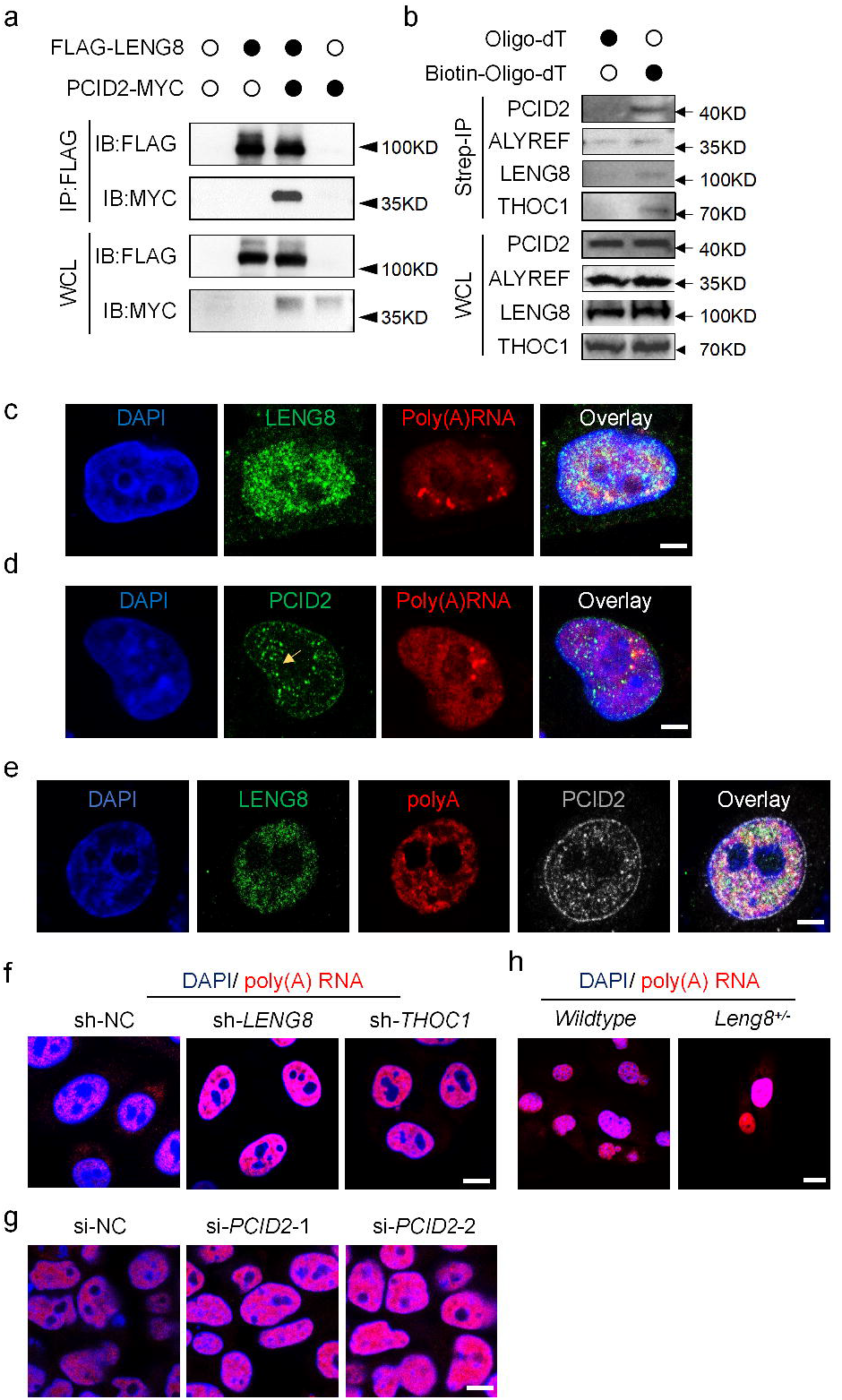
Requirement of LENG8-PCID2 complex in mRNA export. (**a**) HEK293T cells were transfected with FLAG-LENG8 and MYC-PCID2 and then the lysates were sent to immunoprecipitation using anti-FLAG, and Western blot using anti-FLAG and anti-MYC. (**b**) Hela cell lysates were sent to streptavidin-affinity purification using biotin-labeled or non-biotin oligo-dT (25), and western blot using anti-PCID2, anti-THOC1, anti-ALYREF or anti-LENG8. (**c**,**d**) Hela cells were sent to in situ hybridization using Cy3-labelled oligo-dT and immuno-fluorescence staining using anti-LENG8(c) and anti-PCID2(d). (**e**) Hela cells were transfected with 3 x FLAG, and sent to in situ hybridization using Cy3-labelled oligo-dT and immuno-fluorescence staining using mouse anti-FLAG and rabbit anti-PCID. (**f**)Hela cells stably expressing shRNA targeting *LENG8* or *THOC1* were sent to in situ hybridization using Cy3-labelled oligo-dT. (**g**) Mouse tail fibroblasts were isolated from wildtype or *Leng8*^*+/-*^ mice, and then sent to in situ hybridization using Cy3-labelled oligo-dT. (**h**) Hela cells were transfected with a pool of small interferencing RNA(siRNA)s targeting *PCID2* were sent to in situ hybridization using Cy3-labelled oligo-dT. Unprocessed scans of western blot analysis are available in **Supplementary Figure 10**. Bar = 2 μM in **c**,**d**,**e** and 5μM in **f, g, h**. Data are representative of at least three independent experiments (**a, b, c, d, e, f, g, h**).

To determine a functional role for *LENG8* in the process of mRNA export, we created Hela cells stably expressing shRNAs targeting *THOC1*, or *LENG8*. Our FISH result demonstrated that in *THOC1* deficient, or *LENG8* deficient cells, the cytosolic mRNPs were markedly decreased, that is, the nuclear mRNPs were aberrantly accumulated (**Fig. 2f**). Next, we generated *Leng8* knockout mice using CRISPR-Cas9 genome editing methodology (**Wang et al., 2013**) and isolated tail fibroblasts from WT and *Leng8*^+/-^ mice. Consistently, *Leng8*^+/-^ fibroblasts exhibited greater accumulation of poly(A)^+^ mRNPs in the nucleus when compared to the wildtype cells (**Fig. 2g**). Furthermore, Hela cells transfected with a pool of small interfering RNAs targeting *PCID2*, exhibited aberrant accumulation of poly(A)^+^ mRNA in the nucleus (**Fig. 2h**). Taken together, these results suggest that *LENG8* and *PCID2* are evolutionarily conserved mRNA processing factors capable of regulating the export of mRNPs in mammalian cells.

### LENG8 associates with mRNA processing factors

To identify the proteins associated with LENG8, we generated a cell line from human Hela cells, stably transfected with ZZ-LENG8-FLAG (**Sun et al., 2008**). We then purified the tagged LENG8 from cell extracts, by sequential affinity chromatography. The flagged peptide eluates were then electrophoresed on a 4–12% gradient SDS polyacrylamide gel and visualized using silver staining. The bands of interest were excised and analyzed by mass spectrometry(MS) (**Fig. 3a and Supplementary Table 2**).

**Figure 3.**
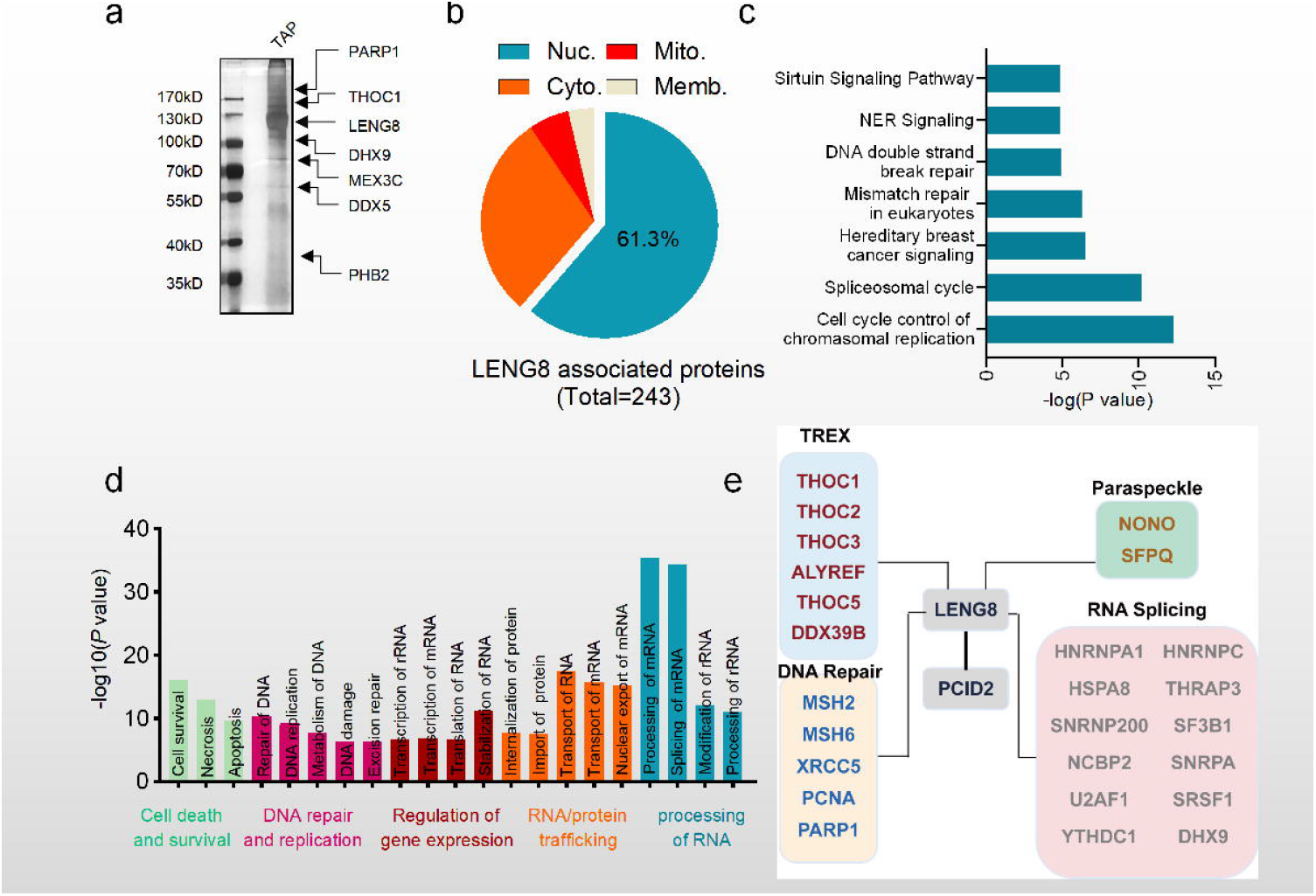
LENG8 associates with mRNA processing factors. (**a**)Silver staining of LENG8 associated proteins by tandem affinity purification. (**b**)Subcellular localization of LENG8 associated proteins. (**c-e**) Molecular function category (**c**), enriched signal pathways (**d**) and protein interaction network (**e**) of LENG8 associated proteins analyzed by Ingenuity Pathway Analysis. Source data of **a** & **c**are in **Supplementary Table 2 & 3**. Data are representative of three independent experiments (**a**).

Our tandem affinity-MS results found that LENG8 significantly enriched for 243 proteins, many of which localized to the nucleus (**Fig. 3b**), or were related to mRNA processing, particularly mRNA splicing and trafficking (**Fig. 3c & 3d, Supplementary Table 3**). As well as PCID2, LENG8 prominently interacted with components of the TREX/THO complex, including THOC1, THOC2, ALYREF, DDX39B, which is consistent with findings from yeast, where thp3 associated with the TREX/THO complex. Furthermore, LENG8 also associated with numerous components of the spliceosome (for example, SNRNP200, SNRPA, U2AF1, SRSF1, and HNRNPA1) and proteins related to the spliceosome cycle, as determined by IPA pathway analysis. In addition, the LENG8 precipitates also contained the paraspeckle components SFPQ and NONO (**Bond et al., 2009**; **Benegiamo et al., 2018**) and several previously identified pre-mRNA processing factors (for example pre-mRNA capping factors NCBP2, THRAP3, and RNA helicase DHX9), as well as DNA repair factors (**Fig. 3e & Supplementary Fig. 4**). Because of the crucial function of the PCI domain as scaffold for protein complex assembly, LENG8 most likely functions as a platform to facilitate multi-step pre-mRNA processing.

We next determined a potential association between LENG8 and the THO complex in mammalian cells. FLAG-tagged LENG8 was co-expressed with HA-tagged THOC (THOC1/3/5/6/7) in HEK293T cells, and a coIP experiment was performed to detect the interaction between LENG8 and THO components. Forty-eight hours after transfection, cell lysates were isolated and subjected to coIP, using anti-FLAG M2 beads. From this, Western blot results revealed that THOC1, THOC5 and THOC6, but not THOC3 or THOC7, were detected in the pellet fraction of FLAG-LENG8 (**Fig. 4a**). We then repeated the coIP assay using HA-LENG8 and FLAG-THOC1, or THOC5, and the resulting pallets were incubated with RNase. Results from the subsequent Western blot, demonstrated that RNase treatment dramatically attenuated the interaction between LENG8 and THOC1, or THOC5 (**Fig. 4b & 4c**), suggesting that these associations are RNA dependent. Immunofluorescence microscopy, further revealed that both ectopically and endogenously expressed LENG8, did indeed co-localize with THOC1, ALYREF and THOC5 (**Fig. 4d-f**). Interestingly, depletion of LENG8 in Hela cells dramatically decreased colocalization of PCID2 with poly(A)^+^ RNAs, but had no effect on the colocalization of THOC1-poly(A)^+^ RNAs **(Supplementary Fig. 5)**, suggesting that the LENG8-PICD2 complex, and THOC1 may exert their functions at different points along the mRNA processing pathway. Taken together, these results suggest that LENG8 is a novel THO complex-associated protein.

**Figure 4.**
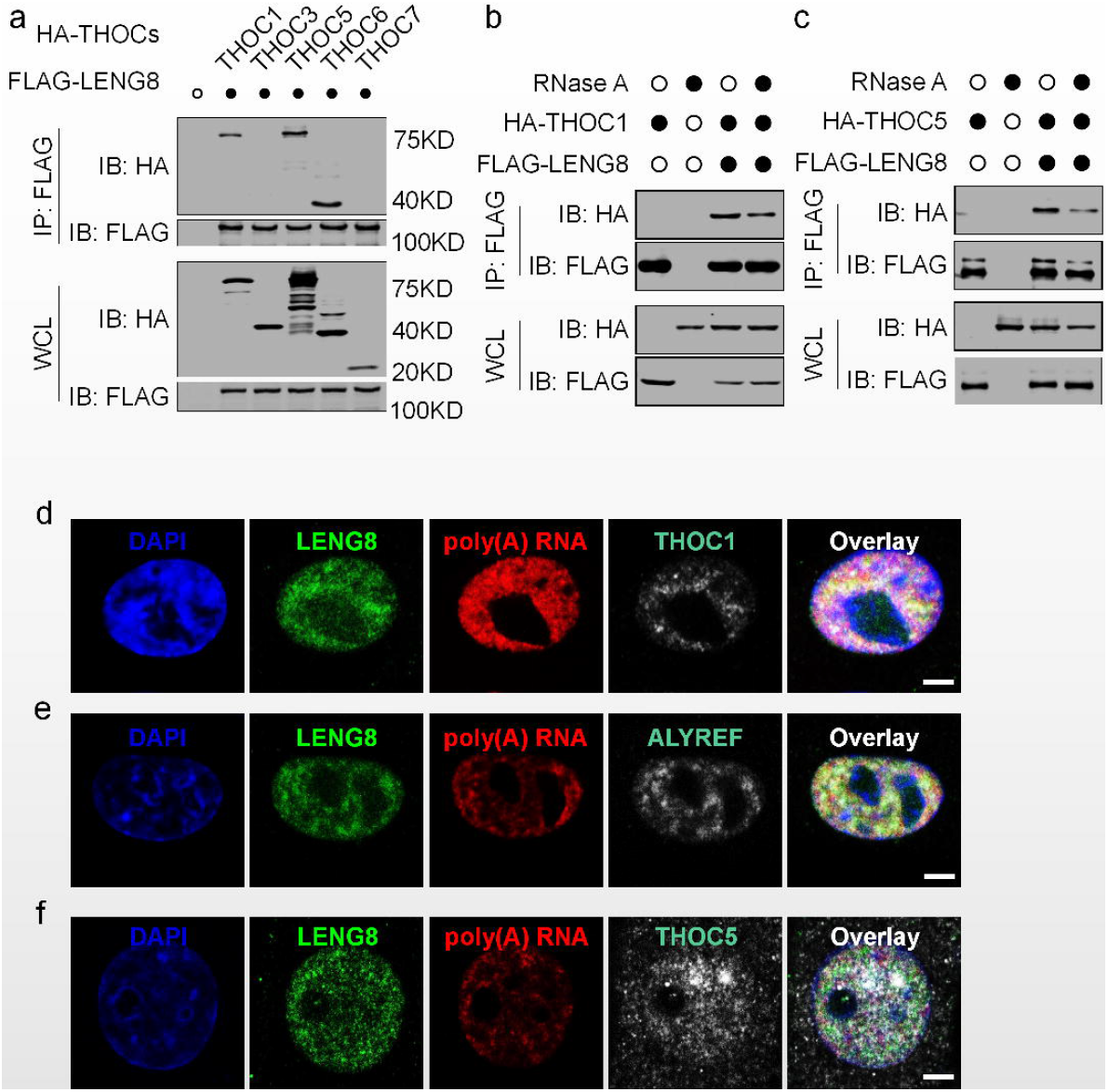
LENG8 associates with TREX components. (**a**) HEK293T cells were transfected with FLAG-LENG8 and indicated plasmids of HA tagged THO components and then the lysates were sent to immunoprecipitation using anti-FLAG, and western blot analysis using anti-FLAG and anti-HA. (**b, c**). HEK293T cells were transfected with FLAG-LENG8 and HA tagged THOC1 (**b**) or THOC5 (**c**) and then the lysates were sent to immunoprecipitation using anti-FLAG. The precipitates were treated with RNAase I and then sent to western blot analysis using anti-FLAG and anti-HA. (**d**,**e**,**f**) Hela cells transfected with 3xFLAG-LENG8 were sent to immuno-fluorescence staining using anti-THOC1 (**d**), ALYREF(**e**) or THOC5 (**f**) and in situ hybridization using Cy3-labelled oligo-dT. Unprocessed scans of western blot analysis are available in **Supplementary Figure 10**. Bar = 2 μM in **d**,**e**,**f**. Data are representative of at least three independent experiments (**a, b, c, d, e, f**).

### RNAs bound by LENG8 and THOC1

Given that LENG8 is associated with the THO complex and is required for mRNA processing, we went on to identify specific RNAs that bind LENG8 or THOC1. Therefore, we precipitated GFP-tagged THOC1, LENG8 or control GFP and sequenced their associated RNAs, using deep sequencing (RNA immune-precipitation followed by deep sequencing, RIP-Seq). We found that when LENG8 was used as bait, 13,328 transcripts from 5,796 genes, were enriched, whereas THOC1 was associated with 32,542 transcripts from 6,150 genes, and 5,179 transcripts from 2,979 genes were enriched in both (**Fig. 5a**). Next, individually enriched RNAs were plotted and individual RNA species were color coded (**Fig. 5b**). Most of the LENG8 and THOC1-binding sites were located in protein-coding transcripts (mRNAs, 73.5 % and 75.6%, respectively). Whereas non-coding RNAs represented 15.9% and 14.2%, respectively and unprocessed RNAs 10.2% and 9.4%. In contrast, LENG8 and THOC1 were minimally bound to pseudogenes (0.6% and 0.2%, respectively), ncRNA (0.1% and 0.1%, respectively) and TEC (To be Experimentally Confirmed) (0.1% and 0.1%, respectively) (**Fig. 5c**). Metagene profiling indicated that compared to THOC1, LENG8 possessed a greater proportion of binding sites in the 5’ UTR (27.1% *vs* 13.4%) and 3’ UTR (36.9% *vs* 28.2%) of mRNAs, but less in exonic (30.9% *vs* 39.6%) and intronic regions (4.7% *vs* 17.7%) (**Fig. 5d & Supplementary Fig. 6**). Gene ontology analysis indicated that LENG8 preferentially enriched mRNAs encoding proteins primarily located in cytosolic organelles, especially mitochondria, and were involved in mitochondrial membrane assembly, mitochondrial gene translation, and metabolism (**Fig. 5e**). In contrast, proteins encoded by THOC1 enriched mRNAs were preferentially located in chromatin and the nucleosome and were involved in regulation of gene silencing and chromatin function (**Fig. 5f**). Furthermore, HOMER motif analysis showed that LENG8 preferentially bound to the ‘C(U/A)GG(A/U)G’ consensus sequence found in both mRNAs and lncRNAs (**Fig. 5g**), whereas THOC1 preferentially bound the “GUGGA(C/U)” consensus sequence. (**Fig. 5h**). Interestingly a m^6^A recognition motif “UGGAC” was found in the consensus sequence of THOC1 bound transcript(**Fig. 5i**), suggesting THOC1 as a potential m^6^A to regulate mRNP biogenesis. Taken together, these data indicated that LENG8 and THOC1 may have distinct functions and associate with different parts of the mRNA processing pathway.

**Figure 5.**
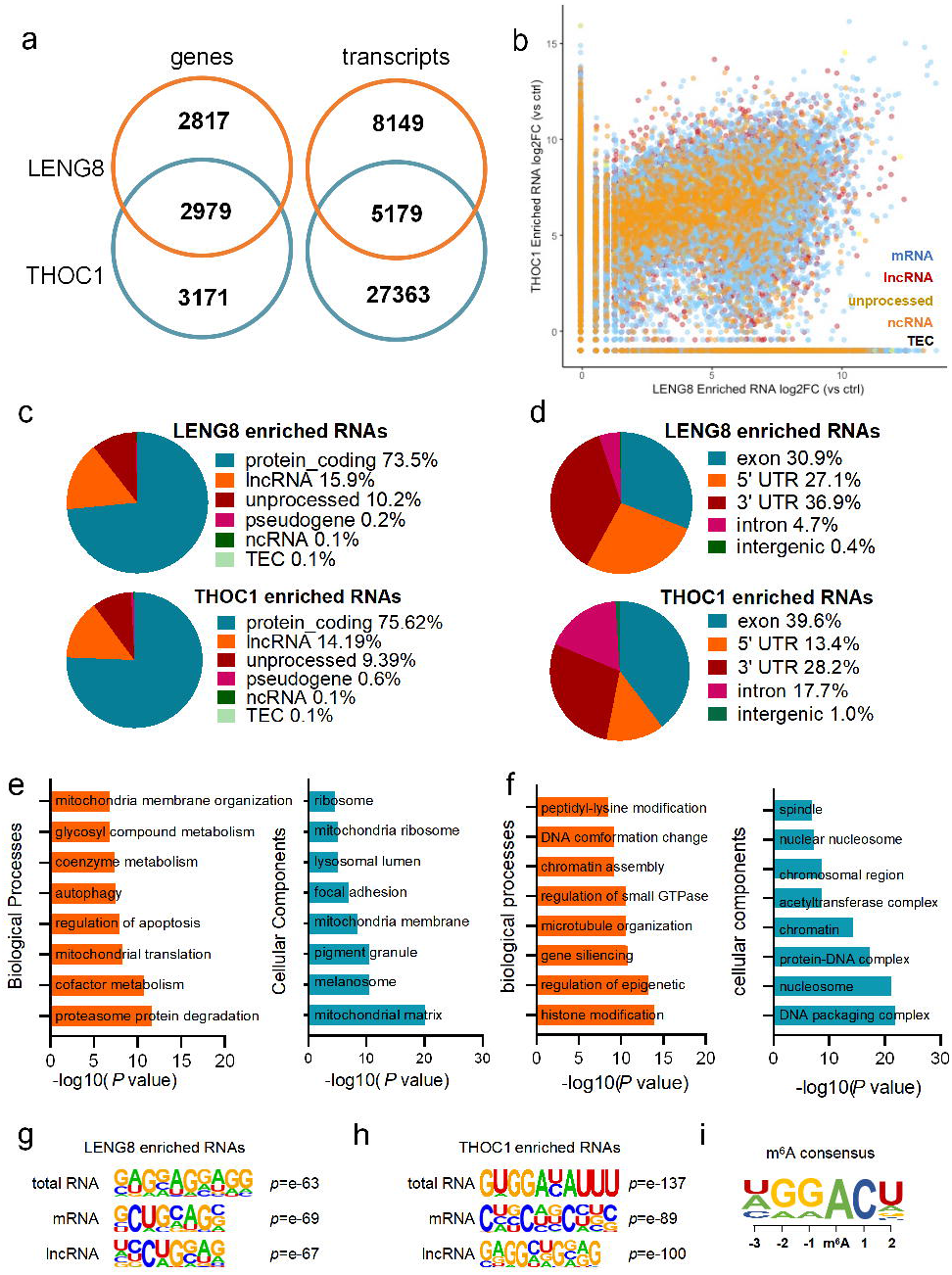
Global analysis of RNAs bound by LENG8 or THOC1. (**a**)Venn diagram showing the numbers of shared high-confidence genes or transcripts enriched by LENG8 and THOC1. (**b**) Scatter plot showing enrichment of transcripts binding to LENG8 (x axis) and THOC1 (y axis) quantified on the gene level. RNAs are colour-coded according to their annotated RNA types in Ensembl. (**c**) Percentage of enriched RNA types binding to LENG8 or THOC1. (**d**)The distribution of LENG8 or THOC1 binding peaks within different gene regions. (**e**,**f**) Gene ontology analysis of LENG8(e) or THOC1(f) enriched RNAs. (**g, h**)Top consensus sequences of LENG8(**g**) or THOC1(**h**) binding sites detected by HOMER Motif analysis.(**i**) m^6^A consensus sequences. Source data of **b** are in **Supplementary Table 4**.

We found that LENG8 and THOC1 bound with high-confidence to *VDAC1* and *VDAC2* mRNA(**Supplementary Fig. 7a**), and this finding was validated using precipitated and quantitative RT-PCR protocols (**Fig. 6a**). To determine which exported mRNAs were influenced by LENG8, we extracted both cytosolic and nuclear mRNAs from Hela cells stably transfected with *LENG8* shRNA and identified them using deep sequencing. Plots of individually sequenced RNAs from both the nuclear and cytosolic fractions after *LENG8* knockdown, revealed that 2,608 transcripts (18.1% of total transcripts sequenced), the nuclear-cytosol export of which were reduced (using a cutoff fold change >2) after *LENG8* depletion. (**Fig. 6b**). However, when the cutoff fold change was reduced to 1.5, *LENG8* knockdown prevented the nuclear export of 8,664 transcripts (59.54% of total transcripts sequenced) (**Supplementary Fig. 7b**). Using IPA signaling pathway analysis, autophagy and mitochondrial damage response pathways were enriched after inhibition of *LENG8* (**Supplementary Fig. 7c**), while Gene Set Enrichment Analysis(GSEA) analysis enriched categories including “Cytokines-cytokine receptor interaction”, “Neuroactive ligand-receptor interaction” and “Olfactory transduction”(**Supplementary Fig. 7d**). Furthermore, *LENG8* deficiency resulted in nuclear retention of 68.5% of those mRNAs bound by LENG8 and 50.8% of those bound by THOC1 in Hela cells (**Fig. 6c**). In addition, the aberrant nuclear accumulation of *VDAC1* and *VDAC2*, two high-confidence targets of *LENG8* and *THOC1*, were validated by quantitative RT-PCR (**Fig. 6d**). FISH results using *VDAC1*-specific probe showed that the cytosolic *VDAC1* mRNA was dramatically reduced after *LENG8* or *PCID2* knockdown(**Fig. 6e**). Therefore, these data, are in strong agreement with our FISH results implicating LENG8 as a necessary factor for mRNA export.

**Figure 6.**
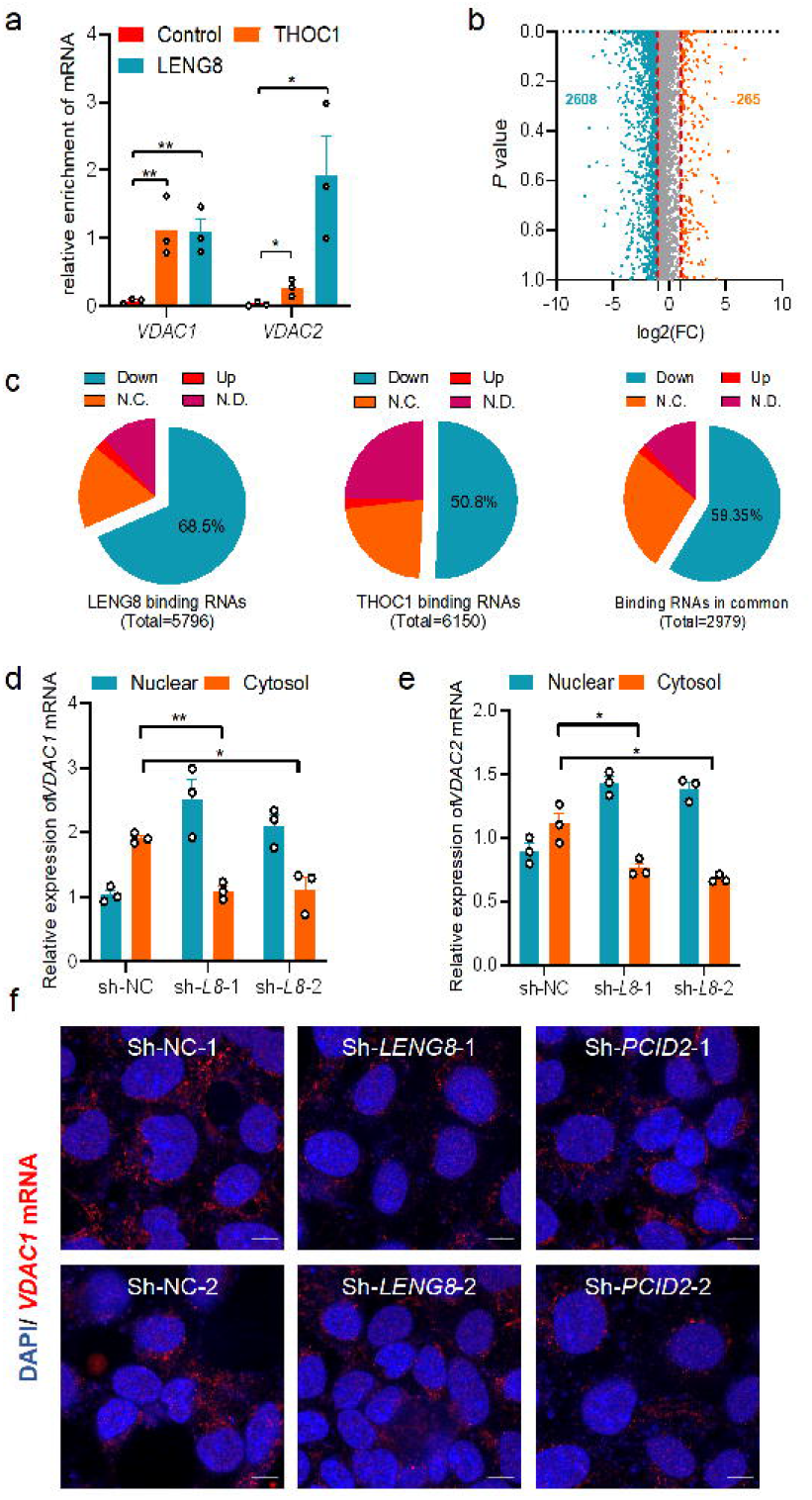
LENG8 is required for *VDAC1*/*2* mRNA export. (**a**)Validation of RIP-sequencing data by qRT-PCR. RNA in LENG8, THOC1 or control (ctrl) precipitates was amplified by qRT-PCR using specific primers for two mRNAs (*VDAC1* and *VDAC2*). (**b**) Hela cells stably expressing shRNA targeting *LENG8* were sent to cytosol-nuclear fractioning, and total RNA from each fraction was extracted and sent to RNA sequencing. Scatter plot indicates individual RNAs sequenced. X axis shows Log2foldchanges of the ratio of Cytosol/Nuclear RNAs after *LENG8* knockdown. (**c**) Analysis of the cytosol/nuclear ratio of mRNA enriched by LENG8, THOC1 or both. (**d**,**e**) Quantitative RT-PCR analysis of cytosol or nuclear *VDAC1*(**d**) and *VDAC2*(**e**) mRNA after *LENG8* knockdown. (**f**) FISH analysis of *VDAC1* mRNA after *LENG8* or *PCID2* knockdown. * p < 0.05 and ** p < 0.01 by the unpaired t-test (**a, d, e**). Data are from three independent experiments with biological duplicates in each (**a, d, e**; mean ± s.e.m. of n=3 duplicates).

### LENG8 controls mitochondrial activity

Given that LENG8 preferentially bound mRNAs encoding mitochondrial proteins, we therefore hypothesized that the activity of *LENG8* was required for maintenance of mitochondrial integrity and activity. To test this, we generated *LENG8* deficient Hela cells using shRNA, or guide RNA-Cas9/CRISPR mediated gene knockout(**Supplementary Fig. 8**) and then examined the morphology and respiration activity of the mitochondria. We determined mitochondrial activity using three different mitochondria-specific labels, capable of detecting mitochondria during respiration (Mitotracker deep red), total mitochondria (Mitotracker green) and ROS-generating mitochondria (MitoSOX) (**Zhou et al., 2011**). The mitochondria from Hela cells with *LENG8* knocked down, or knock out, exhibited reduced ROS production and loss of mitochondrial membrane potential (**Fig. 7a & b**). Similarly, silencing of *THOC1, THOC2* or *ALYREF/THOC4* in these cells, also lead to a reduction of Mitotracker deep red staining, indicating an essential role for the THO complex in the regulation of mitochondrial respiratory activity.

**Figure 7.**
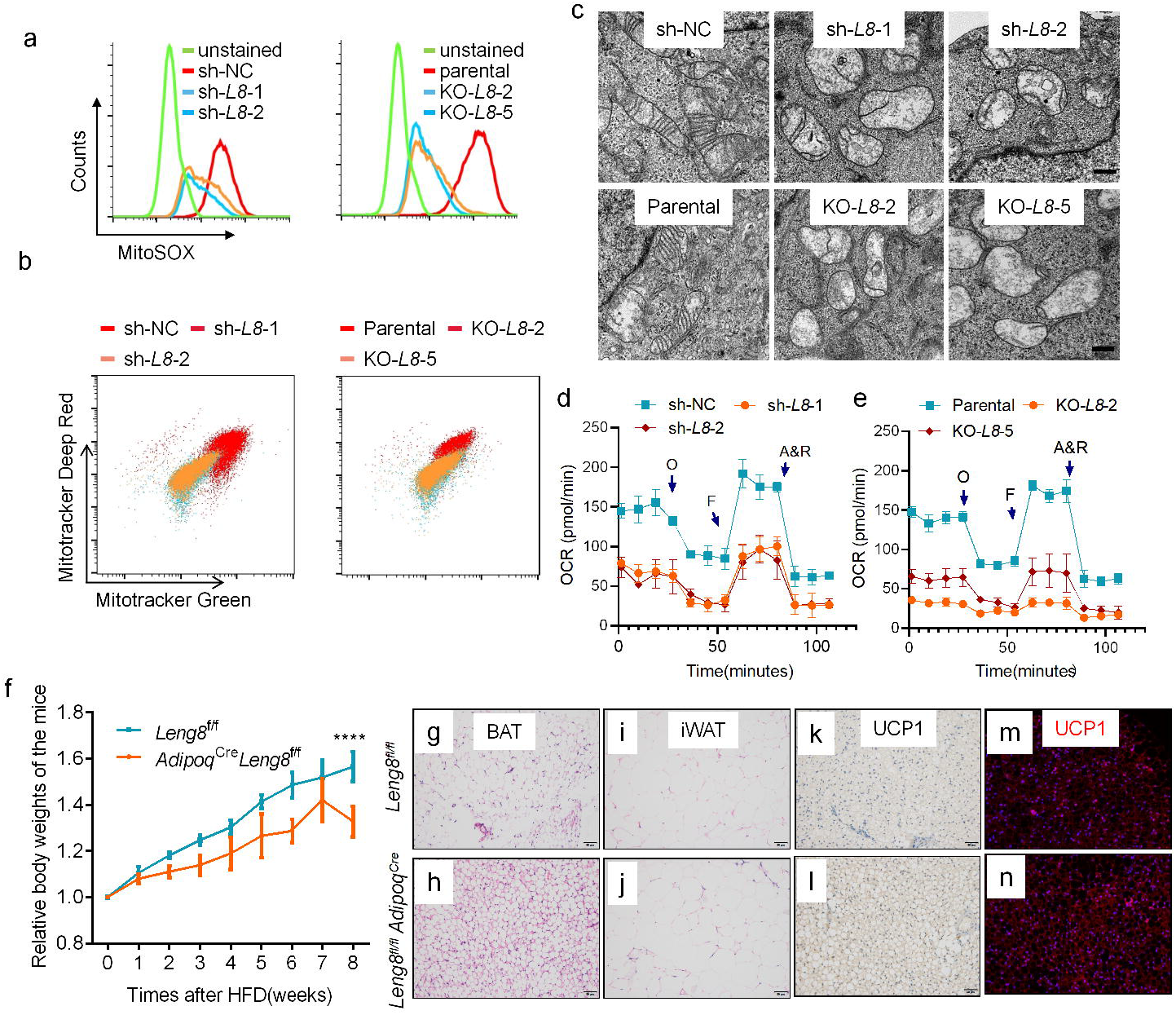
LENG8 controls mitochondria activity. (**a**,**b**) Hela cells of *LENG8* knockdown or knockout were stained with Mitotracker green and Mitotracker deep red (a) or MitoSOX(b) for 30 min and analyzed by flow cytometry.(**c-e**) Hela cells of *LENG8* knockdown or knockout were sent to electron microscopy imaging (**c**) or X24e seahorse test(**d**,**e**). O for oligomycin, F for FCCP, A for antimycin and R for retenone. (**f**) Relative body weight of *Leng8*^*fl/fl*^*Adipoq*^*Cre*^ and *Leng8*^*fl/fl*^ mice after fed with high fat diet. (**g-j**) HE staining of brown adipose tissue(**g, h**) and inguinal white adipose tissue of *Leng8*^*fl/fl*^*Adipoq*^*Cre*^ and *Leng8*^*fl/fl*^ mice after fed with high fat diet(**i, j**). (**k-n**) BAT from *Leng8*^*fl/fl*^*Adipoq*^*Cre*^ and *Leng8*^*fl/fl*^ mice after fed with HFD were sent to immune-staining against UCP1. Data are representative of at least three independent experiments (**a, b, c, d**).

Imaging of Mitotracker green staining using confocal microscopy, found that after application of RNAi directed towards *LENG8*, the cells exhibited enhanced disruption of the mitochondrial network and increased mitochondrial depolarization. (**Supplementary Fig. 9**), further confirming a role for *LENG8* in the maintenance of mitochondrial health. Furthermore, electron microscopy, revealed a complete absence of ‘‘normal’’ mitochondria in *LENG8* deficient Hela cells, including several with a swollen morphology and dramatically decreased numbers of cristae (**Fig. 7c**). In parallel with these morphological changes, *LENG8* deficient Hela cells exhibited a dramatically reduced oxygen consumption rate (OCRs), both in the presence and absence of oligomycin (**Fig. 7d & 7e**).

Mitochondrial activity is tightly associated with thermogenesis in adipose tissue. To explore the physiological function of *LENG8*, we generated the adipose-tissue specific *Leng8* knockout mice by crossing the mice carrying gene-targeted floxed *Leng8* alleles(*Leng8*^*fl/fl*^) with a Cre recombinase (Cre) transgenic mice driven by Adipoq promoter(*Adipoq*^*Cre*^). When fed with high fat diet(HFD), *Leng8*^*fl/fl*^*Adipoq*^*Cre*^ mice exhibited a decreased body weight compared to *Leng8*^*fl/fl*^ mice(**Fig. 7f**). Furthermore, *Leng8* knockout mice displayed marked reductions in adipocyte area and perimeter in sections of inguinal fat pads(iWAT) and brown adipose tissue(BAT)(**Fig. 7g-j**). Consistently, *Leng8* knockout mice displayed marked increase in UCP1 immunolabeling in BAT(**Fig. 7k-n**). Taken together, our finding suggests an essential role for LENG8 in the maintenance of mitochondrial integrity and activity.

## DISCUSSION

Past research employing genetic and biochemical approaches in yeast, have revealed novel regulatory factors involved in mRNA processing, some of which have recently been identified and characterized in mammals (**Katahira and Yoneda, 2009; Yuan et al., 2018; Domínguez-Sánchez et al., 2011; Cheng et al., 2006**). However, it remains unclear as to how this mRNA processing is regulated. The Thp3-csn12 protein complex, has been found to regulate transcriptional elongation and mRNA maturation in yeast, but the existent of mammalian homologues and the specific mechanisms involved in this process remains unknown (**Jimeno et al., 2011; Kragelund et al., 2016)**. Here, we have identified the LENG8-PCID2 complex, as the mammalian equivalent of Thp3-Csn12. LENG8 is a novel regulator of mRNA processing in mammals as well as being multifunctional, as it can bind mRNA associated with mRNA processing machinery and regulating mRNA export. We have also found that LENG8 preferentially binds mRNAs encoding proteins localized to the mitochondria, and its activity is required for the maintenance of mitochondrial morphology and function (**Supplementary Fig. 11**). Importantly, this work has revealed novel aspects of an evolutionarily conserved mRNA processing complex, with the ability to control mitochondrial activity.

In both yeast and mammals, the TREX/THO complex can couple and coordinate transcription, RNA processing and maintain genome integrity (**Domínguez-Sánchez et al., 2011; Gaillard et al., 2007**). Until now, the functional role of LENG8 has remained uncharacterized. However, recently, several interactome studies have revealed resident LENG8 in the RNA polymerase II complex (**Baillat et al., 2005**), RNA processing machinery (**Gebhardt et al., 2015**; **Cano et al., 2015**; **Viita et al., 2019**) and the DNA repair complex (**Alsulami et al., 2019**; **Hu et al., 2019**), suggesting a possible role for *LENG8*, as a master coordinator of these processes. Clearly, to determine the potential ability of *LENG8* to synchronize mRNA processing with genome integrity, warrants further investigation.

Surprisingly, the human mitochondrial genome contains only 13 coding genes, while the number of its nuclear-encoded genes, accounts for 99% of all mitochondrial proteins (**Hendrickson et al., 2010**). It is therefore widely recognized that the expression of these nuclear genes accounts for the control of all aspects of mitochondrial activity, including its morphology, redox regulation, and energetics (**Karakaidos and Rampias, 2020**). Our findings suggest an essential role for *LENG8* in the regulation of mRNA processing involved in mitochondrial activity. To our knowledge, this is the first study involving a link between the maturation of nuclear mRNAs and the regulation of mitochondrial activity. Our results demonstrate that LENG8 can associate with many mitochondrial proteins, including PHB1, a key regulator of mitochondrial homeostasis. PHB1 can respond to mitochondrial stress by translocating from the mitochondria to nuclear regions, where it can regulate the expression of nuclear genes essential for mitochondrial biogenesis, regeneration and degradation (**Hernando-Rodríguez et al., 2018**). Furthermore, these results are consistent with a previous finding demonstrating that LENG8 is also highly enriched in the PHB1 interactome (**Xu et al., 2016**). Taken together, these findings suggest a close association between LENG8 and PHB1. However, a role for PHB1 in the mitochondrial stress response through its association with LENG8 remains to be explored.

N6-methyl-adenosine (m^6^A) is a newly-characterized RNA fate determiner and affects multiple aspects of mRNA processing including splicing, translation and decay through various m^6^A recognition proteins, the so-called m^6^A readers(**Frye, M et al., 2018**). Several studies have shown that the m^6^A reader YTHDC1 recruits TREX complex to mRNA and its deficiency lead to aberrant nuclear accumulation of mRNA(**Roundtree IA et al., 2017; Lesbirel, S et al., 2018**), suggesting an essential role of m^6^A modification for mRNA nuclear export. Here we showed that THOC1 itself, which is the most well-characterized TREX component, is a potential m^6^A reader. Furthermore, YTHDC1, as well as another m^6^A reader YTHDF2, were found to co-precipitate with LENG8 in this study. These finding strengthens the link between m^6^A modification and mRNA export. However, our RIP-seq results do not support LENG8 as a significant m^6^A reader. Thus, whether *LENG8* is involved in m6A mediated mRNA export, as well as other processes of RNA biology, remains to be explored.

We conclude that *LENG8* is a fundamental factor involved in the support of mitochondrial activity. Furthermore, this has been seen previously in *C. elegans*, where *Hpo-10* was shown to be the nematode homologue of *LENG8*/*Thp3* and may be a candidate for the regulation of the mitochondrial unfolded protein response (**Liu et al., 2014**). Furthermore, they also found that *Thoc-1* was able to regulate mitoUPR in their reverse-genetic screening analysis. Collectively, these findings suggest an essential role for TREX/THO-mediated mRNA maturation during the regulation of mitochondrial homeostasis in diverse species.

## Materials and Methods

### Constructs and Antibodies

The nucleotide sequence used for *LENG8, PCID2, THOC1, THOC2, THOC3, THOC5, THOC6, THOC7* over-expression were NM_052925.3, BC016614, BC010381, NM_001081550.1, NM_032361.3, NM_003678.4, NM_001142350.1 and NM_025075.3. All of these constructs were purchased from Sinobiological (Beijing). The mouse anti-LENG8 were produced by Daian Biotechnology (Wuhan, Hubei) using a recombinant LENG8 fragment (1-300 aa). The rabbit anti-PCID2 were from Abcam; the rabbit anti-THOC1, THOC2, THOC5, ALYREF, Myc-tag were from Abclonal (Wuhai, Hubei); the mouse anti-GFP were from Roche; the mouse anti-FLAG was from Genscript (Nanjing, Jiangsu). FITC-conjugated goat anti-mouse, Cy3 and Cy5 conjugated goat anti-rabbit were from Beyotime biotechnology (Shanghai); HRP conjugated goat anti-rabbit and goat anti-mouse were from Abclonal (Wuhan, Hubei). Anti-FLAG agarose beads and streptavidin-affinity magnetic beads were from Genscript (Nanjing, Jiangsu); Protein A/G magnetic beads were from Thermo Fisher Scientific.

### Cell culture and transfection

HEK293T, Hela were obtained from cell bank of Shanghai Institute for Biological Sciences (CAS) and maintained in standard conditions (37°C, 5% CO2) in DMEM/High Glucose medium with 10% FBS, 100 U/ml penicillin and100 mg/ml streptomycin (Life Science). All cell lines have been tested negative for mycoplasma contamination.

Cells were seeded the day before transfection. The next day, when cells were 70%–80% confluent, the medium was changed to penicillin- and streptomycin-free medium. DNA in Opti-MEM (Life Technologies) was mixed with Lipofectamine 2000 or Lipofectamine 3000 (Invitrogen) and then incubated for 20 min at room temperature, then added dropwise to the cells. The medium was changed to complete medium after 6 h and the cells were used in experiments at 48h-72h post-transfection.

### shRNA mediated gene knockdown and gRNA-Cas9 mediated gene editing

For shRNA mediated gene knockdown, more than two effective shRNA clones to each target coding sequence were prepared. shRNA sequences were cloned into the lentiviral expression plasmid pLKO.1 and transfected into HEK293T cells to generate recombinant lentiviruses. Hela or Hep2 cells were transduced with the lentiviral supernatants and selected with 1 mM puromycin (MCE). RT-qPCR and western blot analysis was performed to verify significant depletion of each target sequence. The shRNA sequences used are listed in **Supplementary Table 6**.

The *Leng8*^*-/-*^ mice were generated through the CRISPR/Cas9 method as described previously. Briefly, in vitro-translated Cas9 mRNA and gRNA were co-microinjected into the C57BL/6 zygotes. The pair of gRNA sequences used to generate the knockout mice is GCTATGTGCCACCTTCAGCT and ACTAGGACATGCTAATGTCC. Founders with frame shift mutations were screened by DNA sequencing. One F0 founder, of which 937bp fragment containing exon3 and exon4 of Leng8 loci was deleted, was crossed with C57BL/6 wildtype and 11 F1 mice were got for the *Leng8*^*-/-*^ mice. One of the F1 mice was chosen to backcross to the WT mice more than 10 generations to maintain the strain. All protocols were approved by the local ethics committee of Shanghai JiaoTong University Affiliated Sixth People’s Hospital.

### Immunofluorescence staining and *in situ* hybridization of cultured cells

Cells were cultured on coverslips or glass bottom dishes, fixed with 4% PFA, permeabilized with 0.1% Triton X-100 in PBS and blocked with 1% BSA. For immunofluorescence staining, cells were incubated with primary antibodies at 4°C overnight and then secondary antibody or DAPI at room temperature for 30 min. Antibodies were used at the following dilutions: rabbit polyclonal anti-PCID2, THOC1, THOC5, ALYREF, 1:200; mouse monoclonal anti-LENG8, 1:100; Cy3-conjugated goat anti-rabbit IgG, 1:500; and FITC-conjugated goat anti-mouse IgG, 1:500. For *in situ* hybridization, cells were incubated with 5 μM cy3 labeled oligo-dT (70) in 2 × SSC buffer at 42 °C overnight. Samples were examined and the figures were acquired with an LSM 710 confocal laser-scanning microscope (Carl Zeiss, Oberkochen, Germany) at 63× or 100× magnification.

### RNA-Immunoprecipitation and Sequencing

HeLa cells expressing GFP-tagged human *THOC1* or *LENG8* were pelleted by centrifugation at 500g for 10 min at 4 °C and washed twice with ice-cold PBS. Cells were lysed in an equal volume of RIP lysis buffer (10 mM HEPES pH 7.0, 100 mM KCl, 5 mM MgCl2, 25 mM EDTA, 0.5% (v/v) Nonidet-P40, 1 mM dithiothreitol, protease inhibitor cocktail (EDTA-free, Beyotime) for 30 min on ice in the presence of 100 U ml^−1^ RNase inhibitor (Sangon)) and lysates clarified by centrifugation at 9,000g and 4 °C for 10 min. Clarified lysates were incubated with anti-GFP with a final concentration at 0.2ug ml^-1^ and the binding reactions were conducted for 2 hours at 4 °C with continuous gentle rotation. The reaction mixture was centrifuged at 2000 x g at 4 °C for 5 minutes to remove debris. The ChIP grade protein A/G magnetic beads were washed with RIP binding/wash buffer (50 mM Tris pH 7.4, 150 mM NaCl, 1 mM MgCl2, 0.05% (v/v) Nonidet-P40) containing 25 mM EDTA, protease inhibitors and 100 U ml^−1^ RNase inhibitor for 3 times. The mixture of RNA-protein complexes was added to 40 μl of 100 % beads, and then the binding was conducted overnight at 4 °C on a rotary wheel, followed by five washes with RIP binding/wash buffer. Beads were then resuspended in Trizol (Invitrogen) and RNA was isolated according to the manufacturer’s instructions. Two RNA immunoprecipitations per bait were carried out in parallel. RNA quality was assessed on a Genetic Analyzer (Agilent) and TruSeq RNA library construction and next-generation sequencing were performed by the Lianchuan Biotechnology (Hangzhou, Zhejiang). All samples were sequenced on an Illumina HiSeq2500 platform at 15 million 100-bp single reads per sample. After quality control of the sequencing libraries, reads were trimmed and mapped against the Ensembl genome annotation and the human genome assembly (hg19/GRCh38) using Tophat2. Reads mapping to ribosomal RNAs or the mitochondrial genome were removed. RNAs binding to THOC1 or LENG8 were identified by differential quantification (bait over control) against the Ensembl genome annotation using cuffdiff from the cufflinks package. RNAs with fold changes >2, FDR corrected P values <0.01 and minimal read counts of 10 were considered as enriched. To discover preferences of THOC1 and LENG8 for different RNA species, we extracted RNA types and gene-model-related features from the Ensembl annotations and plotted them using custom scripts.

### Affinity purifications

For affinity purifications with biotin-labelled mRNA, Hela cells were pelleted and lysed in an equal volume of AP lysis buffer (10 mM HEPES pH 7.0, 150 mM KCl, 5 mM MgCl2, 25 mM EDTA, 0.5% (v/v) Nonidet-P40, 1 mM dithiothreitol, protease inhibitor cocktail (EDTA-free, Beyotime) for 30 min on ice in the presence of 100 U ml^−1^ RNase inhibitor (Sangon)) and lysates clarified by centrifugation at 9,000g and 4 °C for 10 min. Clarified lysates were incubated with biotin conjugated oligo-dT(25) or non-biotin conjugated control oligo-dT with a final concentration at 5 μM and the binding reactions were conducted for 2 hours at 4 °C with continuous gentle rotation. The reaction mixture was centrifuged at 2000 x g at 4 °C for 5 minutes to remove debris. Streptavidin-affinity magnetic beads were washed with AP binding/wash buffer (50 mM Tris pH 7.4, 100 mM NaCl, 1 mM MgCl2, 0.02% (v/v) Nonidet-P40) containing 25 mM EDTA, protease inhibitors and 100 U ml^−1^ RNase inhibitor for 3 times. The mixture of RNA-protein complexes was added to 20 μl of 100 % beads, and then the binding was conducted overnight at 4 °C on a rotary wheel, followed by five washes with AP binding/wash buffer. The RNA-protein coated beads were boiled in 50 μl 1 x SDS loading buffer, and subjected to SDS–polyacrylamide gel electrophoresis and western blot analysis.

### Tandem affinity purification of LENG8 complex

The tandem affinity purification strategy to fractionate the LENG8 complexes from Hela cells was performed as follows. The stable cell line capable of expressing ZZ-LENG8–3xFLAG was obtained. Thus, the cells were grown in DMEM with 10% FBS plus 1% P/S and harvested near confluence. The cell pellet was washed with chilled PBS three times and then lysed in an equal volume of TAP lysis buffer (50 mM Tris-Cl pH 7.4, 100 mM KCl, 5 mM MgCl2, 25 mM EDTA, 0.5% (v/v) Nonidet-P40, 1 mM dithiothreitol, protease inhibitor cocktail (EDTA-free, Beyotime) for 30 min on ice. The homogenate was centrifuged for 20 min at 10,000 x g. The supernatant was transferred to a fresh tube. Then 50 μl of packed IgG beads was added to the 4 mg protein extract, followed by gentle rotation overnight at 4 °C and then washed by TAP lysis buffer of 50 mM KCl for 3 times. The bound protein was eluted by TEV protease cleavage and further purified by anti-FLAG -conjugated beads. The final eluates from the FLAG beads with FLAG peptide were resolved by SDS/PAGE on a 4–12% gradient gel and visualized by silver staining. Specific bands were cut off and subjected to mass spectrometry analysis. The protein interaction network analysis was performed using QIAGEN’S Ingenuity Pathways Analysis (QIAGEN’S Ingenuity pathway analysis, Ingenuity Systems, http://www.qiagen.com/ingenuity, version 52912811).

### Nucleocytoplasmic separation, RNA isolation and sequencing

Cells were washed with cold PBS, and then incubated at -20 °C for 5 minutes. Subsequently, buffer A containing 10 mM HEPES (pH 7.9), 1.5 mM MgCl2, 10 mM KCl, 0.5mM dithiothretol (DTT), and 1 mM PMSF was added. The cytoplasmic and nuclear fractions were separated by centrifugation at 17,000 × g for 30 minutes at 4 °C. Both of the cytoplasmic and nuclear fractions were then resuspended in Trizol (Invitrogen) and RNA was isolated according to the manufacturer’s instructions. All samples were sequenced on an Illumina HiSeq2500 platform at 15 million 100-bp single reads per sample. After quality control of the sequencing libraries, reads were trimmed and mapped against the Ensembl genome annotation and the human genome assembly (hg19/GRCh38) using Tophat2. Reads mapping to ribosomal RNAs or the mitochondrial genome were removed.

### Structure modelling for LENG8 and THP3

The structure modeling of LENG8 and THP3 protein were conducted by Swiss-model (https://swissmodel.expasy.org/)(**Waterhouse A *et al*., 2008**). The template used for this modeling is Sac3:Thp1:Sem1 complex (PDB Accession: **3t5v.1**), and all figures were generated using Chimera V1.14.

### Western blot

Proteins were lysed from cells using RIPA buffer containing 10 mM Tris-Cl, pH 8.0, 150 mM NaCl, 1% Triton X-100, 1% Na-deoxycholate, 1 mM EDTA, 0.05% SDS and fresh 1× proteinase inhibitor. The protein concentration was determined via the Bradford method using the Bio-Rad protein assay before proteins were equally loaded and separated in polyacrylamide gels. The proteins were then transferred to a nitrocellulose filter membrane (Millipore) and were incubated overnight with indicated primary antibodies. HRP-conjugated secondary antibodies were then applied to the membrane, and the Western blot signal was detected using auto-radiographic film after incubation with ECL (GE Healthcare) or SuperSignal West Dura reagents (Thermo Scientific).

### Q-PCR

For detection of *LENG8* and *PCID2*, total RNA from Hela cells was extracted with TRIzol reagent according to the manufacturer’s instructions (Invitrogen). One microgram of total RNA was reverse transcribed using the ReverTra Ace® qPCR RT Kit (Toyobo, FSQ-101) according to the manufacturer’s instructions. A SYBR RT-PCR kit (Toyobo, QPK-212) was used for quantitative real-time PCR analysis. The relative mRNA expression of different genes was calculated by comparison with the control gene Gapdh (encoding GAPDH) using the 2-△△Ct method. The sequences of primers for the qPCR analysis are shown in **Supplementary Table 6**.

### Mitochondrial activity studies

For OCAR measurement, an XFe24 Extracellular Flux analyzer (Agilent) was used to determine the bioenergetic profile of wildtype and *LENG8* deficient Hela cells. Hela cells were plated at 1,000,000 cells per well in XFe24 plates 24 before the Mito Stress Tests. All assays were performed following manufacturer’s protocols. Results were normalized to cell number.

For mitochondrial mass measurement, wildtype and *LENG8* deficient Hela cells were incubated with Mitotracker green and Mitotracker deep red at 50nM for 30 min at 37 °C. Mitochondria-associated ROS levels were measured by staining cells with MitoSOX at 2.5 μM for 30 min at 37 °C. Mitochondria membrane potential was measured using the kit from Invitrogen and performed according to the manufacturer’s instructions. Cells were then washed with PBS solution and resuspended in cold PBS solution containing 1% FBS for FACS analysis.

### Statistical analysis

The results are represented as the mean ± s.e.m., and statistical significance between groups was determined using an unpaired t-test or the Mann-Whitney U test. GraphPad Prism software 8.0 was used for all analyses, and a *p<0.05 was considered statistically significant.

## ACKNOWLEDGEMENT

This study was supported by National Science Foundation (No. 81971240 to Liu F); China Postdoctoral Science Foundation (No. 2020M671248 and No.2020T130672 to Zhao YX), Shanghai Yangfan Talents Program (20YF1456200 to Zhao YX), Shanghai Municipal Commission of Science and Technology(No.18DZ2260200), Jinan Science and Technology Development Program (No. 201907018 to Zhang N) and Shandong Provincial Key Research and Development Program (No.2017G006037, No. ZR2020MH201 to Zhang N)

## AUTHOR CONTRIBUTION

Y.X.Z. X.T.W. H.L.Y. and F.L. designed experiments. Y.X.Z. X.T.W., Y.N.L., N.N.L., S.M.W., N.Z., Z.F.G, W.Y.X., L.F.M, C.Y.L., J.G. and Z.L.C. performed experiments and analyzed data. Y.X.Z., Q.R.D. and F.L. prepared the figures and wrote the manuscript.

## DATA AVAILABILITY

Data have been deposited in the Gene Expression Omnibus under accession code GSE171126. Other data that support the findings of this study are available from the corresponding author upon request.

## COMPETING FINANCIAL INTERESTS

The authors declare no competing financial interests.

## Notes

### Competing Interest Statement

The authors have declared no competing interest.

