## Supplementary Files for "LENG8 regulation of mRNA processing, is responsible for the control of mitochondrial activity"

**Supplementary Tables**

**Table 1.** Identities and similarities of TREX components from yeast and human.

**Table 2.** Proteins associated with LENG8 from Affinity-MS identification.

**Table 3.** Molecular functions of LENG associated proteins.

**Table 4.** GO and KEGG annotations of LENG8 or THOC1 enriched RNAs.

**Table 5.** Signaling Pathway analysis of nuclear detained RNAs after LENG8 RNAi in Hela cells by IPA.

**Table 6.** Sequence for primers, shRNAs, siRNAS and gRNAs used.

**Supplementary Figure Legend**

**Figure 1 Alignment of human PCID2, yeast THP1 and CSN12 proteins**

**Figure 2 Alignment of human LENG8, yeast THP3 and SAC3 proteins.**

**Figure 3 LENG8 was a nuclear protein.** (**a**) Hela were transfected with DsRed-*LENG8* and then sent inmunostaining using anti-Nucleolin. (**b**)Hela cells were transfected with 3×FLAG-*LENG8* and then sent to cytosol-nuclear fractioning, and then western blot analysis using anti-FLAG. ME:Membrane extract; CE:Cytosol extract; NE: Nuclear extract; CB：Chromatin bound; PE: Pallet extract. (**c,d**) GFP tagged full-length or indicated truncation of *LENG8* (**c**) were over-expressed in Hela cells and sent to confocal microscopy imaging(**d**).

**Figure 4 Interactome analysis of LENG8 associated proteins by IPA.**

**Figure 5 Colocalization of PCID2 or THOC1 with poly(A) RNA after LENG8 depletion.** Hela cells were transfected with a pool of *LENG8* siRNAs and then sent to in situ hybridization using Cy3-labelled oligo-dT and immuno-fluorescence staining using anti-THOC1 and anti-PCID2.

**Figure 6 Metagene profiles of enrichment of LENG8 and THOC1 binding sites across mRNA transcriptome.**

**Figure 7 LENG8 depletion inhibited mRNA export. (a)** Hela cells stably expressing shRNA targeting LENG8 were sent to cytosol-nuclear fractioning, and total RNA from each fraction was extracted and sent to RNA sequencing. Scatter plot indicates individual RNAs sequenced. X axis shows Log2foldchanges of the ratio of Cytosol/Nuclear RNAs after *LENG8* knockdown. Cut off = +/- 0.58(Fold change=1.5). (**b，c**) enriched signal pathways analysis (**b**) and GSEA analysis(**c**) of genes whose exporting are attenuated after *LENG8* depletion. Source data of **b** are in **Supplementary Table 5**.

**Figure 8 Inhibition of *LENG8* expression by shRNA-mediated knockdown or Cas9-crispr mediated knockout.** (**a**) Total RNAs from Hela cells stably expressing shRNAs targeting control vector or *LENG8* were subjected to quantitative realtime RT-PCR analysis of *LENG8*. (**b**) WT or *LENG8* Knockdown Hela cells were were subjected to Western blot analysis using anti-LENG8. (**c**) Schematic illustrating a single nucleotide deletion in LENG8 exon2 created by Cas9/crispr, and successful deletion was verified by Sanger sequencing as shown. (**d**) Single clones from *LENG8* knockout Hela cells created by Cas9/crispr method were sent to immunoblot validation using anti-LENG8.

**Figure 9 Inhibition of THO complex leads to reduction of mitochondria activity. (a)** Hela cells of indicated genotype were stained with Mitotracker deep red for 30 min and analyzed by flow cytometry. (**b**) Hela stably expressed *LENG8* shRNAs were transfected with mitoGFP, and then sent to confocal microscopy imaging.

**Figure 10 Unprocessed scans of western blot analysis**.

**Supplementary Figure 1**

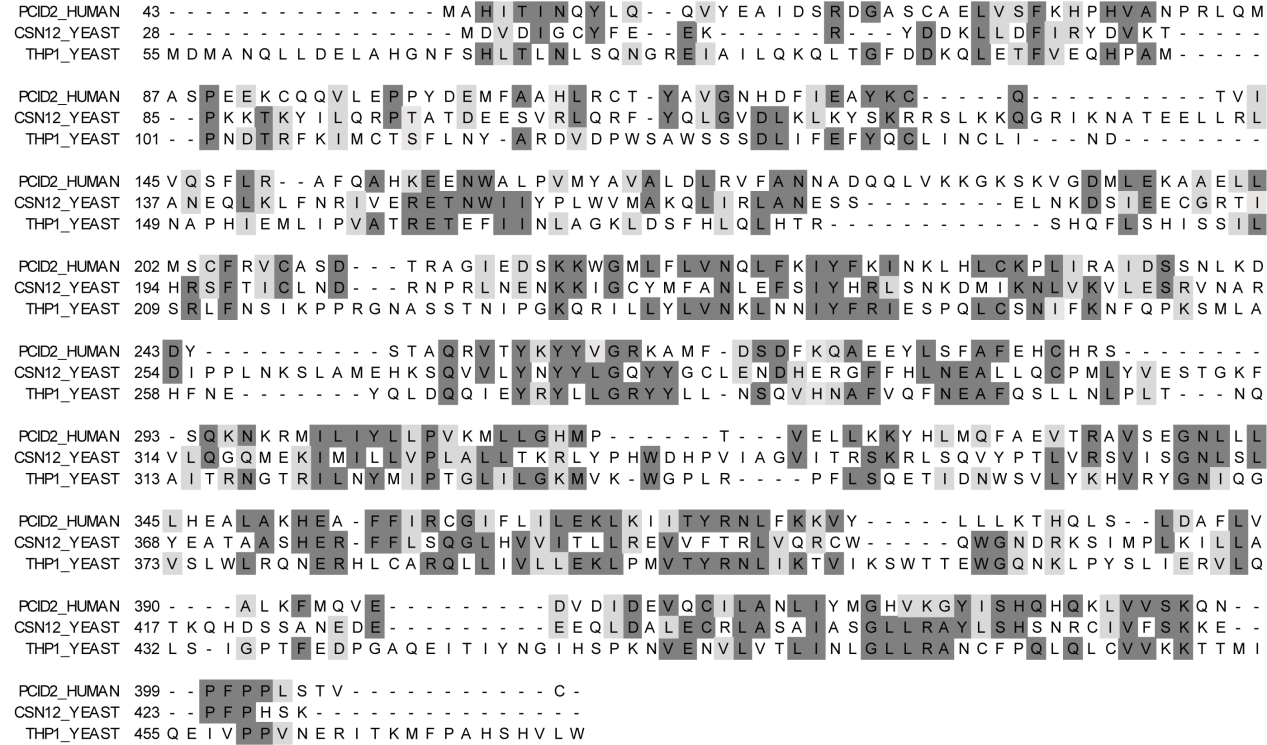

**Supplementary Figure 2**

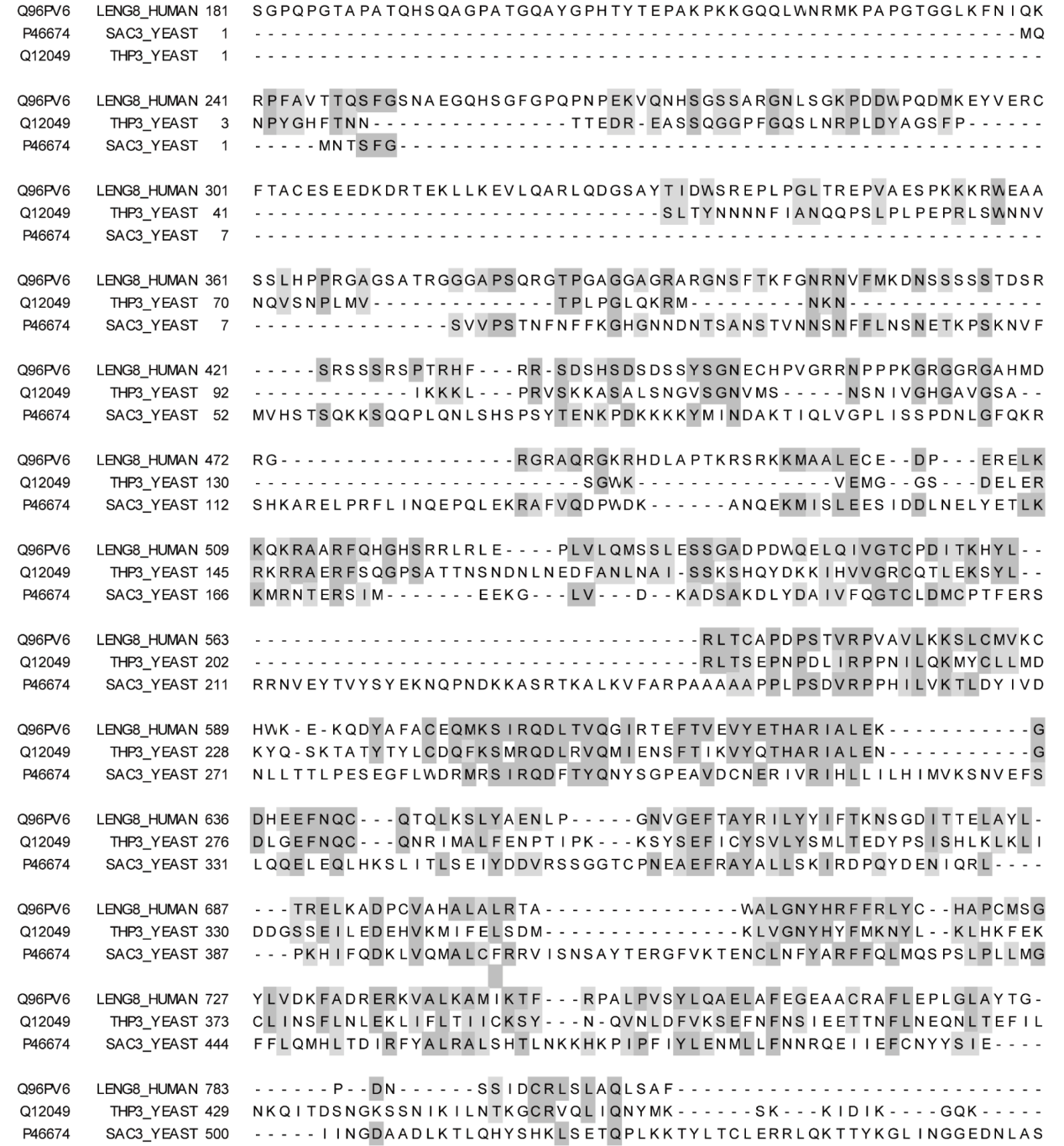

**Supplementary Figure 3**

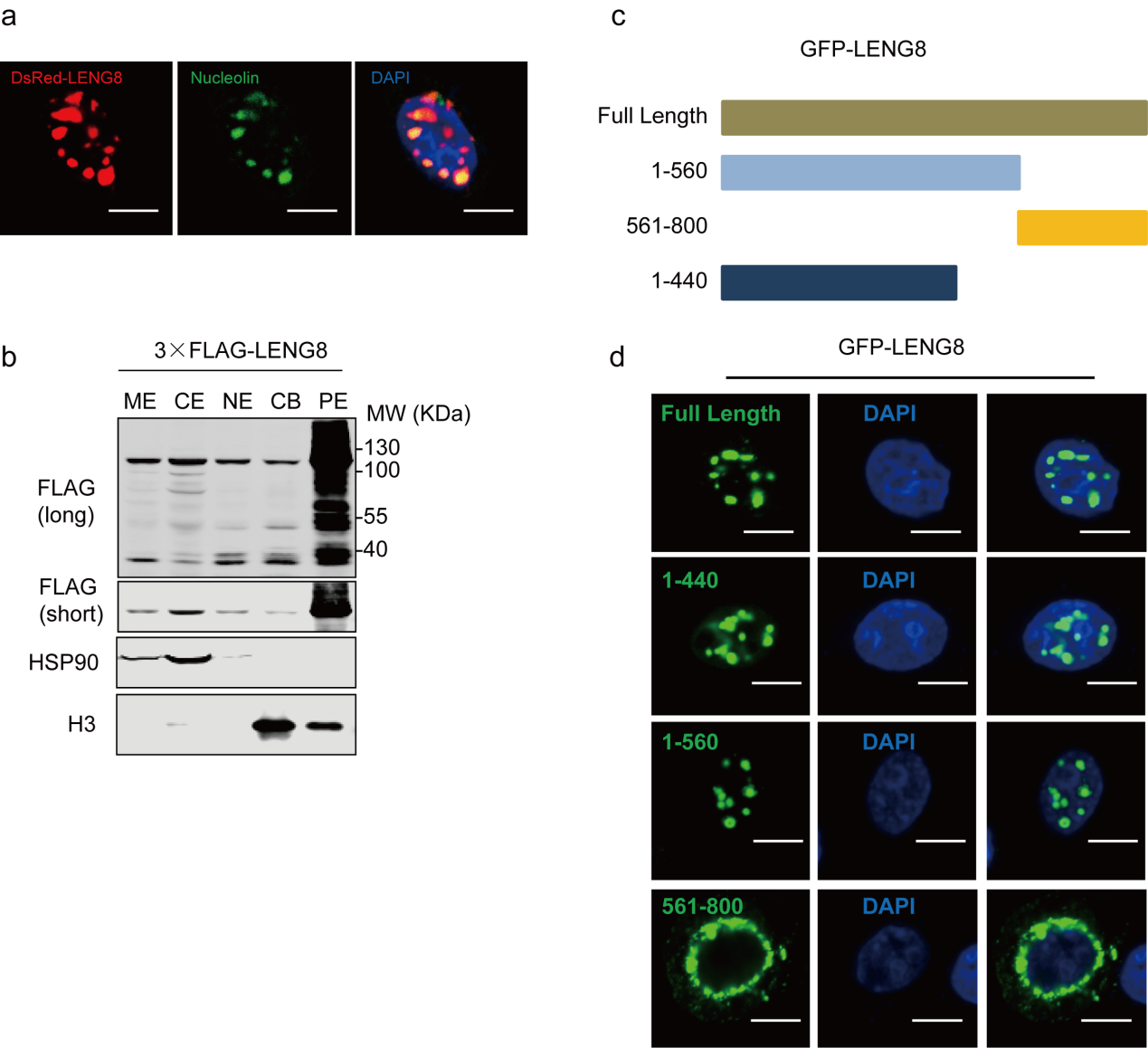

**Supplementary Figure 4**

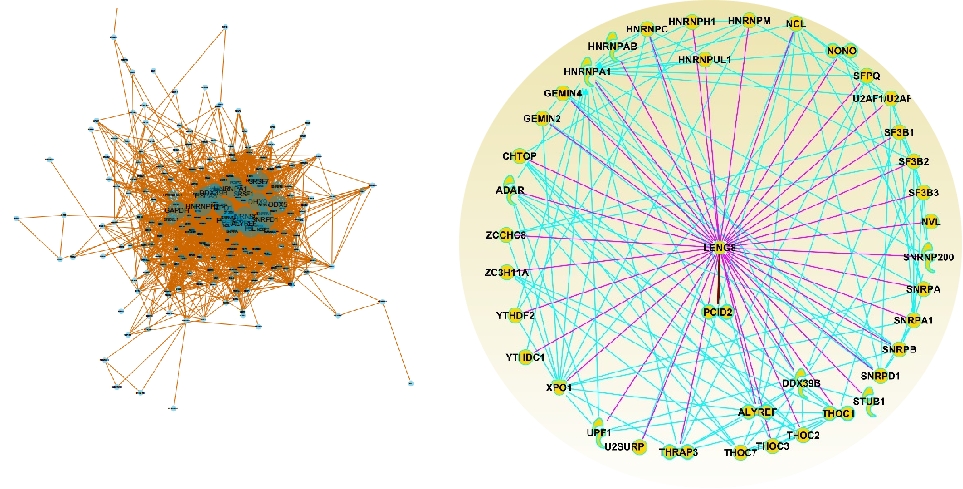

**Supplementary Figure 5**

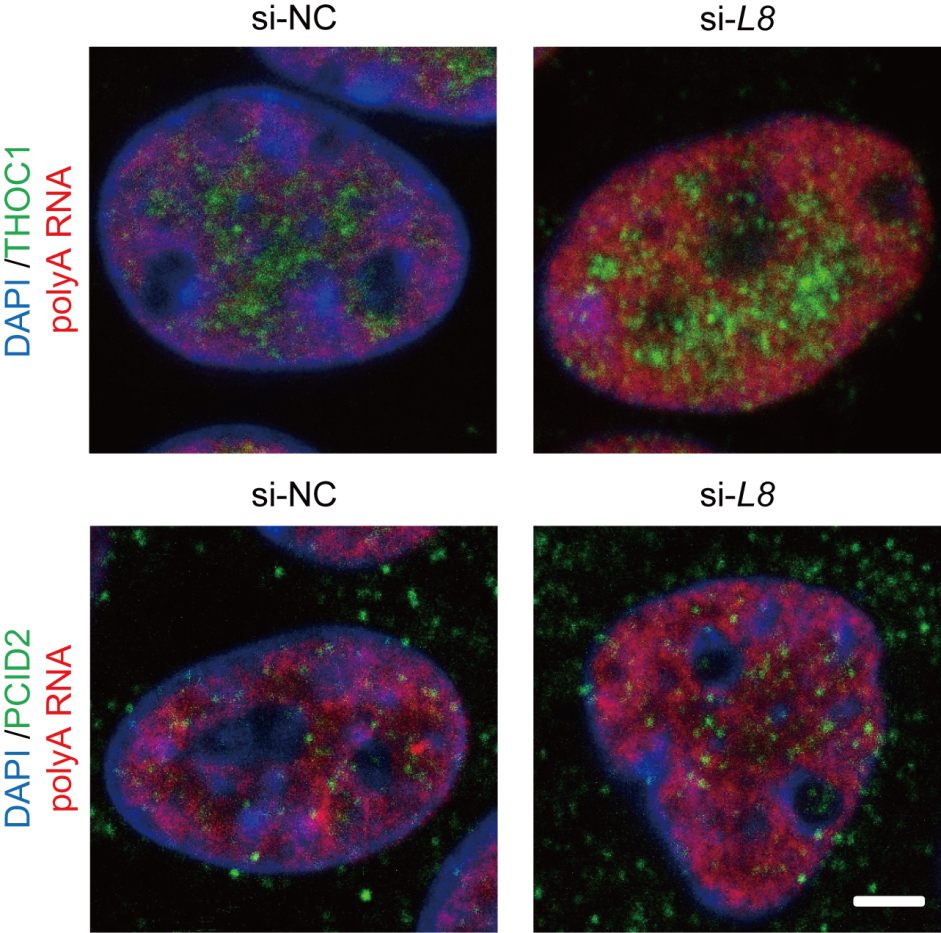

**Supplementary Figure 6**

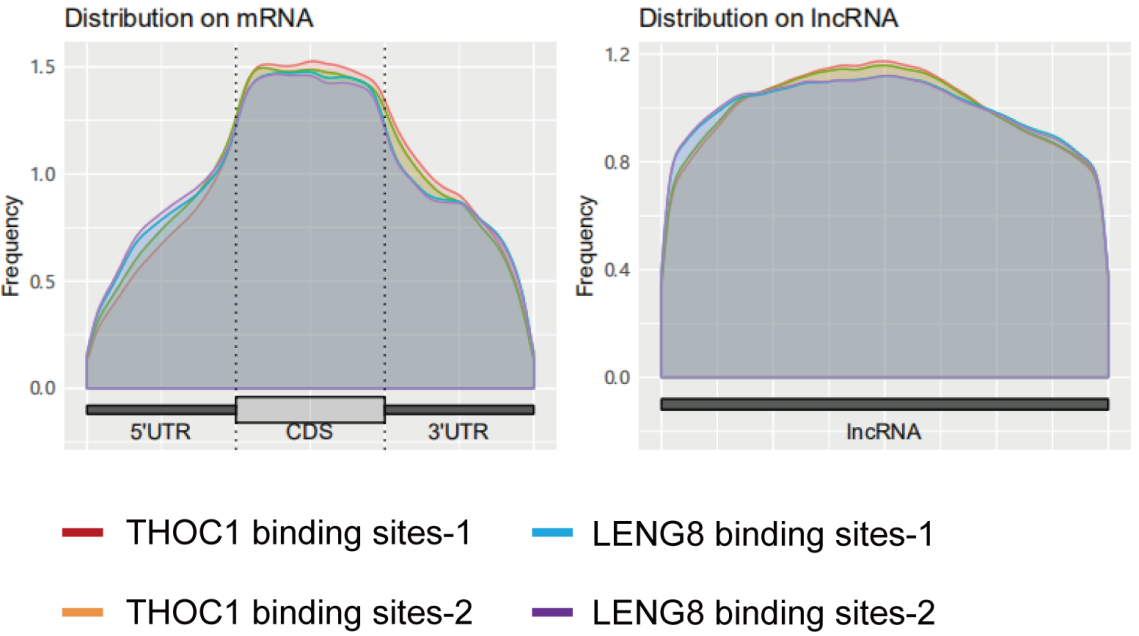

**Supplementary Figure 7**

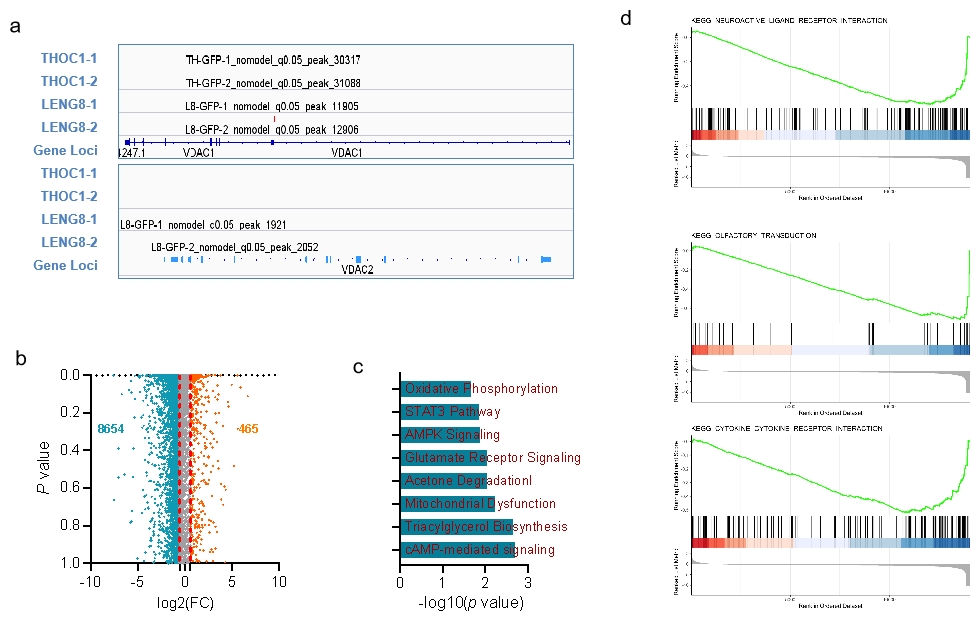

**Supplementary Figure 8**

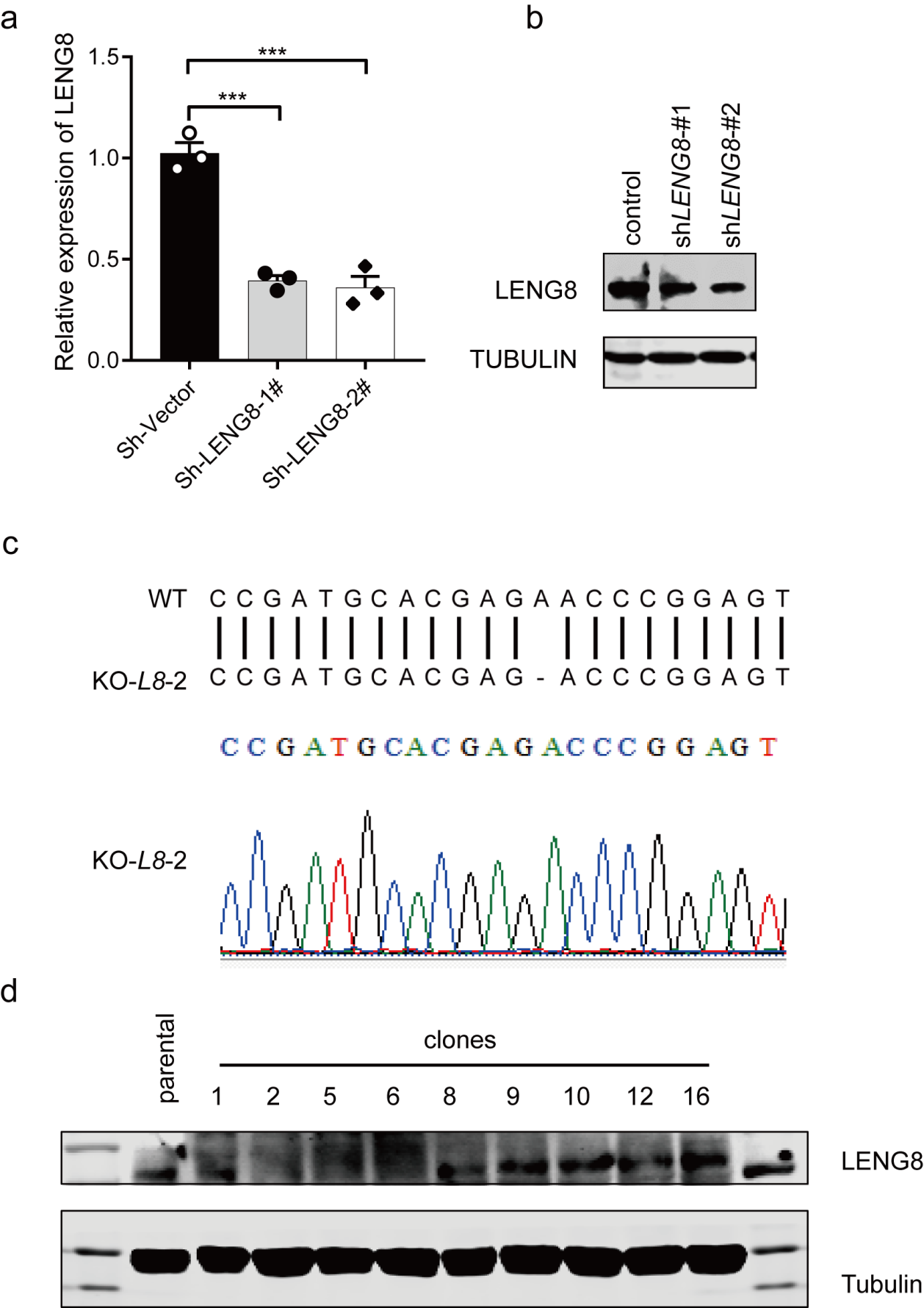

**Supplementary Figure 9**

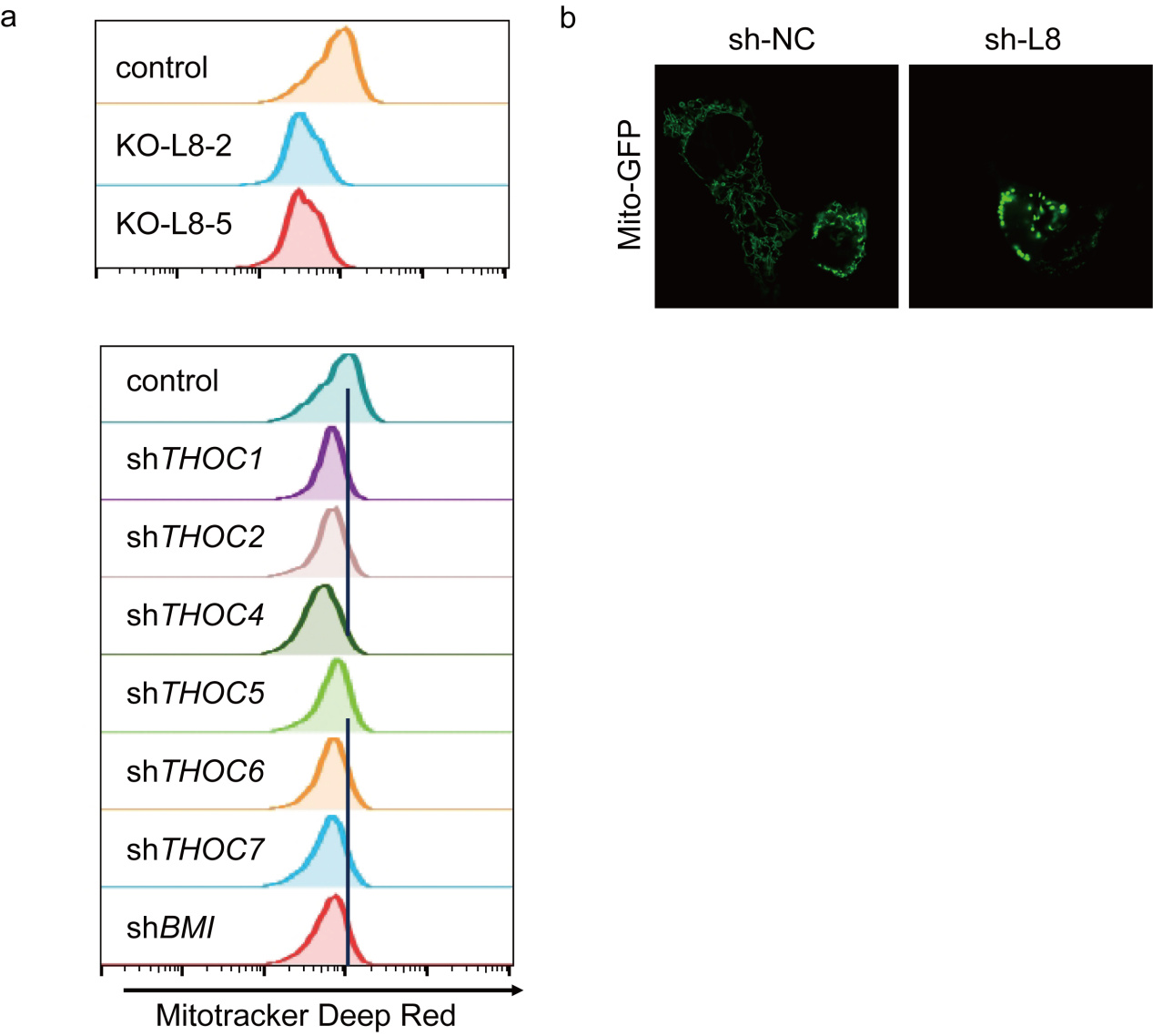

**Supplementary Figure 10**

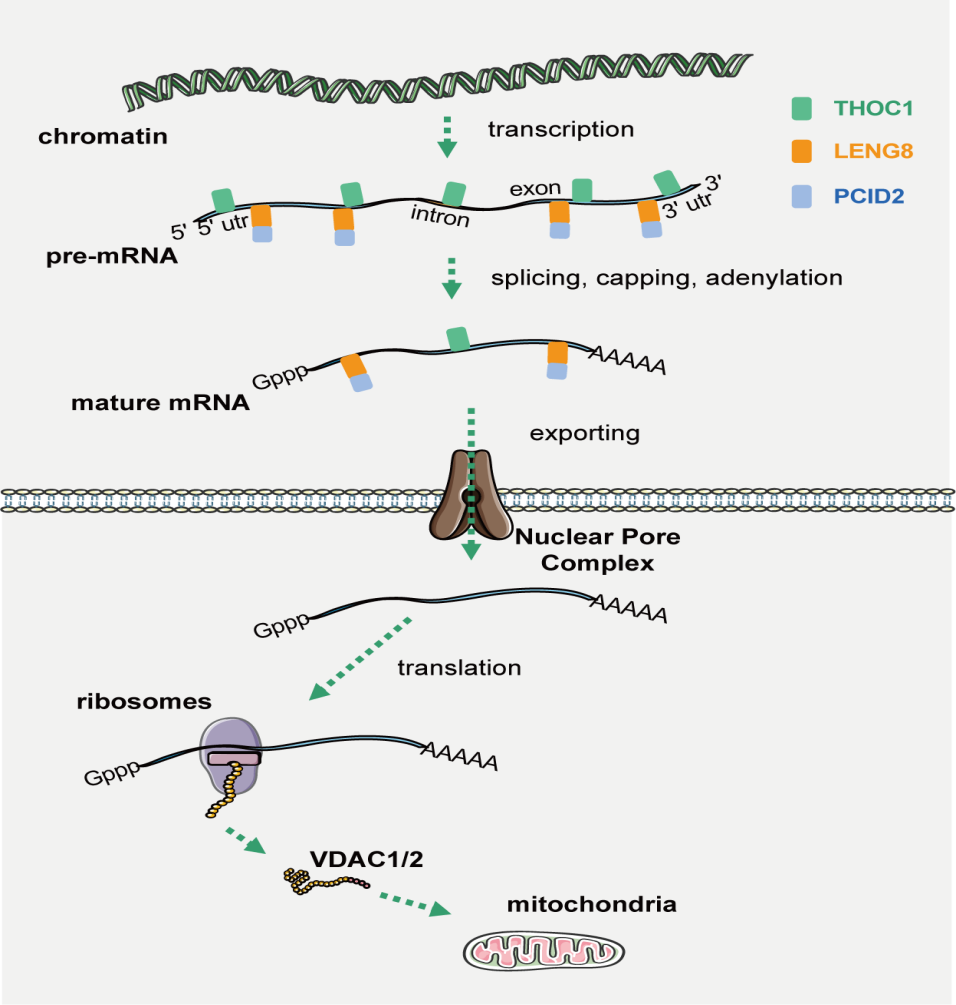

**Supplementary Figure 11**

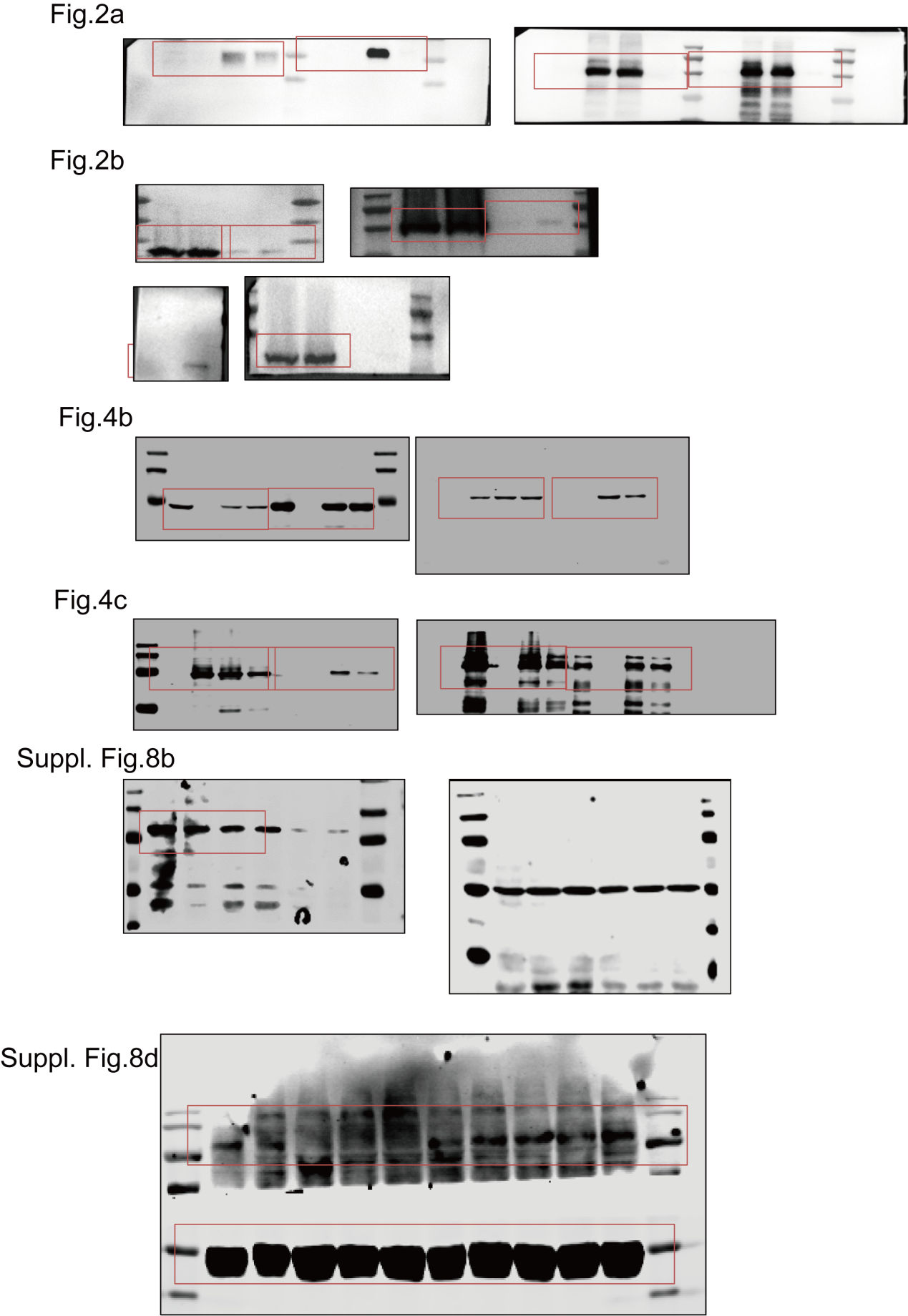
